# Genetic exchange drives relatedness of large linear and circular extrachromosomal elements of archaea

**DOI:** 10.64898/2026.10.02.756104

**Authors:** Natasha J. Jablonsky, Ling-Dong Shi, Daniela S. Aliaga Goltsman, Shuai Wang, Luis E. Valentin Alvarado, Rohan Sachdeva, Gavin J. Knott, Jill F. Banfield

## Abstract

**Background:** Archaeal extra-chromosomal elements (ECEs) play key roles in archaeal host biology and evolution, yet little is known about their diversity and interactions with each other. Here, we report 42 complete 66 - 165 kbp genomes: 23 “Klingons”, 8 “Romulans” and “Romulan-like elements”, and 11 “Ferengi”, three new families of archaeal ECEs from diverse terrestrial environments.

**Results:** Methylation patterns, phylogenetics and gene sharing analyses suggest that Klingons and Romulans associate with *Candidatus* Methanoperedens and Ferengi are likely hosted by *Candidatus* Bathyarchaeaota. Like known archaeal viruses and plasmids, these ECEs have circular genomes, unlike the linear Borg and mini-Borg ECEs that also replicate in *Methanoperedens,* and they replicate bi-directionally. The new ECE genomes all encode genes for methylation, glycosylation, protein degradation, and novel TnpB-like enzymes, which may interact with novel CRISPR-like arrays. All genomes also encode proteins with strong structural similarity to archaeal viral capsid proteins, some of which are predicted to multimerize into pentameric or hexameric subunit capsid-like arrays and thus may encapsulate themselves. The three new ECE families share gene content with each other and with Borgs and mini-Borgs, often concentrated in specific genomic regions. Discordance among single gene and genomic phylogenies suggests that apparent relatedness of the five distinct ECE types stems from extensive horizontal gene transfer, rather than evolution from a common ancestor.

**Conclusions:** The three new families of ECEs broaden our understanding of the diversity of archaeal ECEs and highlight the role of horizontal gene transfer in modulating the archaeal and archaeal ECE gene pools.

## Background

Extrachromosomal elements (ECEs), including viruses and plasmids, can contribute to the metabolic capacity of associated organisms, impact host fitness, and play roles in host genomic evolution, yet archaeal ECEs are greatly understudied relative to their bacterial counterparts. This knowledge gap is detrimental, as archaea can be key members of microbial communities and vital players in global biogeochemical cycles. Archaea play important roles in production and anaerobic oxidation of methane (CH_4_), a potent greenhouse gas, by methanogens and anaerobic methanotrophic “ANME” archaea respectively.

Many ECEs associated with the anaerobic methanotroph *Candidatus* Methanoperedens (formerly ANME-2d) have been reported, including circular plasmids (1), mini-chromosome-like elements (2), mobile genetic elements (3) and viruses (4,5). One family of *Methanoperedens* ECEs was reported to have linear genomes of up to 1.1 Mbp in length, significantly larger than the largest archaeal virus ever reported, the 144 kb *Halogranum* tailed virus 1 genome (6,7). As they could not be classified as plasmids or viruses (8), and in view of their propensity for assimilation of genes from organisms, they were named Borgs (a reference to alien species of the Star Trek series). Borgs encode capsid-like proteins (9) that are predicted to multimerize into capsid-like arrays and share genomic characteristics with Nucleocytoviricota large DNA viruses (10). Subsequently, mini-Borgs were reported (5) and suggested to be ancestral to the Borgs (11), broadening the known size range of this linear ECE family to 49 kb – 1.1 Mb (8).

A prior study reported four unclassified ECEs that were inferred to associate with *Methanoperedens* (5). We sought additional examples of these elements and in this study, report 45 new circular genomes from metagenomes, 42 of which were curated to completion. The genomes range from 53 to 165 kb in length, and all encode many proteins closely related to those of Borgs. We performed phylogenetic and comparative genomic analyses to explore the ECEs’ relatedness with each other and with Borgs and mini-Borgs, analysed the coding structure and evidence for replication modes, and investigated the protein inventories via *in silico* structure prediction. These analyses assigned the ECEs into three families and demonstrated the close relatedness of the new circular ECEs with each other and with linear Borgs and mini-Borgs, due largely to horizontal gene transfer.

## Results

### Groups of novel, large, archaeal ECEs found across environments

We used protein sequences from unclassified elements reported by Shi et al. (5) to identify and reconstruct six related ECEs from metagenomic data from the same wetland soil where they were originally identified, in Lake County, California. The sequencing read archive (SRA) and assemblies of a subset of the SRA (Logan) were also searched and 39 related ECEs were identified. 42 of the 45 genomes were manually curated to completion. In the case of two ECE genomes, one from the Lake County wetland and one from Germany groundwater, assemblies were verified by long-read PacBio sequencing.

The 45 ECEs were recovered from 20 different sampling sites in the US, Canada, Germany, England, China and Australia. Listed by decreasing incidence, the sample types where the novel ECES were derived from include groundwater, wetland soil, terrestrial soil, lake sediment, landfill and peat soil. Six out of the 20 metagenomes contained more than one ECE genome **(Table 1)**. Using read mapping-based coverage data we estimated that DNA from ECEs comprised between 0.42% and 18.51% of the composition of the DNA pool from the 50 most abundant organisms (excluding the ECEs) **(Figure 1)**. The environment with the largest proportion of total ECE DNA is a California wetland sample (SRVP 1.05m, abundance = 18.5%). The environment with the most abundant single ECE was a Chinese groundwater sample (SRR17498766, abundance = 13.12%). The top 5 organisms (excluding ECEs) in groundwater samples and terrestrial soil samples were more frequently bacterial (43/50 and 10/15 respectively), whereas across the wetland samples, the top 5 contigs were all archaeal (20/20), although all five samples were taken from the same location.

**Figure 1:**
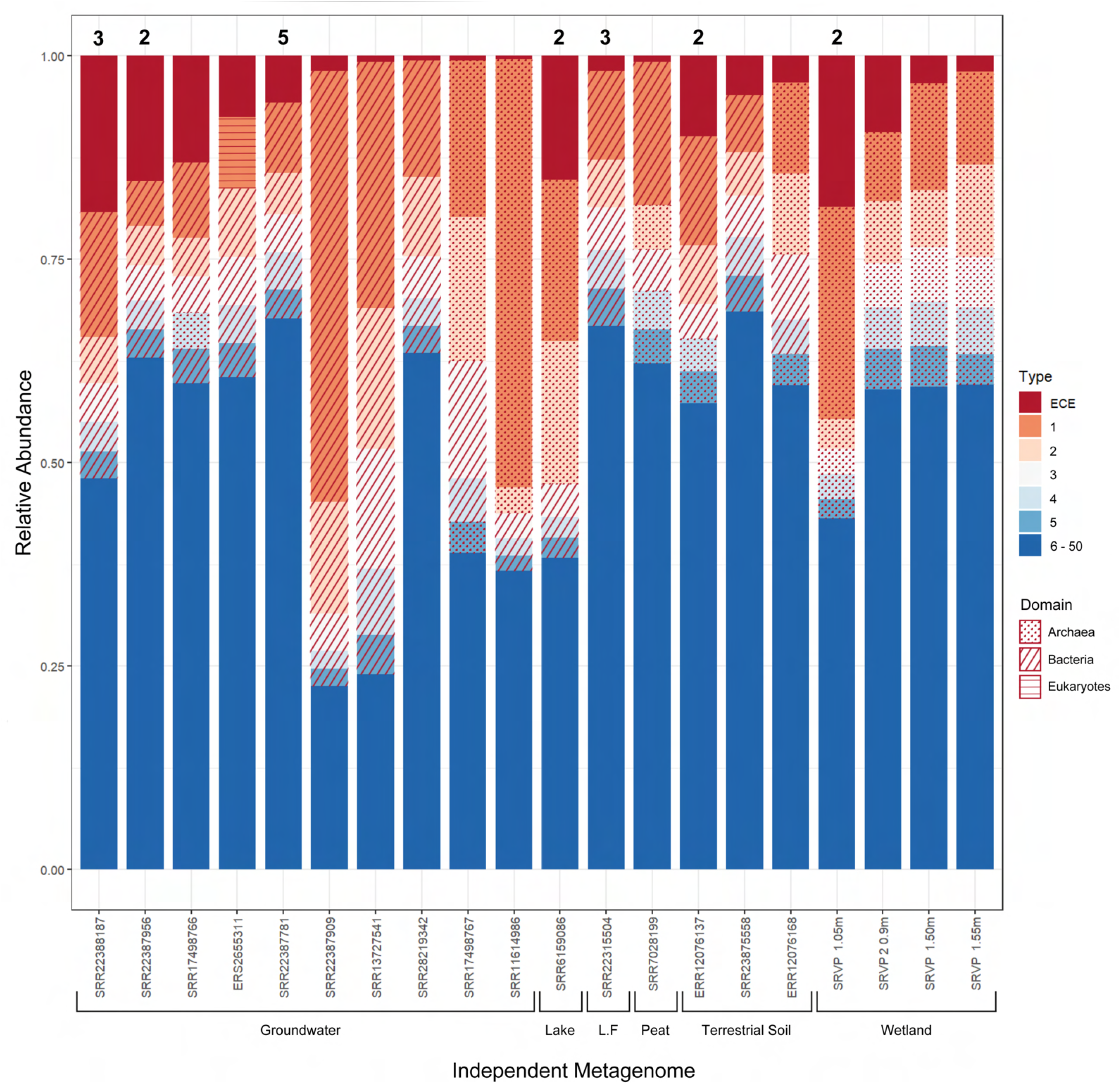
Relative abundance of ECE compared to top 50 contigs from metagenome, grouped according to environment type. Organisms with top five highest coverage values (excluding ECE) are indicated and marked according to domain. “Lake” = Lake Sediment, L.F = Landfill. In cases where more than one ECE was curated from the same metagenome, the number of ECEs is labelled on top of the column.

**Table 1:** Sample site details of curated ECE genomes published in this study.

| SRA/ENA Accession | Location | Coordinates | Depth (m) | Metagenome Type | Genomes in Sample |
| --- | --- | --- | --- | --- | --- |
| **Public Data Samples (from SRA)** |  |  |  |  |  |
| ERR12076137 | Skipton, North Yorkshire, UK | 54.03 N, 2.38 W | 0.5 | Terrestrial | Dargh + Qeng Klingon |
| ERR12076168 | Skipton, North Yorkshire, UK | 54.03 N, 2.38 W | 0.5 | Terrestrial | Jup Klingon |
| ERS2655311 | Little Forest Legacy Site, NSW, Australia | 34.04 S, 150.98 E | 1 | Groundwater | Meq Klingon |
| SRR11614986 | Islinger Muhlbach, Regensburg, Germany | 48.99 N, 12.13 E | 35 | Groundwater | Hov Klingon |
| SRR13727541 | Chochrane, Alberta, Canada | 51.29 N, 114.42 W | 46 | Groundwater | Genorex Romulan |
| SRR17498766 | Hetao Basin, Inner Mongolia Province, China | 40.97 N, 107.01 E | 16 | Sediment | Doq Klingon |
| SRR17498767 | Hetao Basin, Inner Mongolia Province, China | 40.97 N, 107.01 E | 14 | Sediment | Soq Klingon |
| SRR22315504 | Hollandsville, Delaware, USA | 38.96 N, 75.67 W | 3 | Landfill | Chal, Qes + Yiv Klingon |
| SRR22387764 | Changchun, Jilin Province, China | 44.17 N, 125.82 E | 29.6 | Groundwater | Euro Ferengi |
| SRR22387775 | Baicheng, Jilin Province, China | 45.95 N, 123.56 E | 33 | Groundwater | Yuan Ferengi |
| SRR22387781 | Baicheng, Jilin Province, China | 45.62 N, 124.04 E | 30 | Groundwater | Pound, Ruble, Dong, Ringgit + Rupiah Ferengi |
| SRR22387909 | Quanzhou, Fujian Province, China | 24.94 N, 118.52 E | 26.4 | Groundwater | Bas Klingon |
| SRR22387926 | Dalian, Liaoning Province, China | 39.87 N, 123.08 E | 34.5 | Groundwater | Devoras Klingon |
| SRR22387956 | Yiyang, Hunan Province, China | 29.10 N, 112.31 E | 67.06 | Groundwater | Belak + Shaenor Romulan |
| SRR22388041 | Weihai, Shandong Province, China | 37.53 N, 122.07 E | 10.6 | Groundwater | Per Klingon, Rand Ferengi |
| SRR22388071 | Guilin, Guangxi Province, China | 25.28 N, 110.25 E | 70 | Groundwater | Ramjep Klingon |
| SRR22388187 | XiAn, Shanxi Province, China | 34.15 N, 108.40 E | 70 | Groundwater | Nagh Klingon, Dollar + Gold Ferengi |
| SRR22388334 | Fushun, Liaoning Province, China | 41.89 N, 124.011 E | 32.5 | Groundwater | Menthe Klingon |
| SRR23875558 | Shanghai, China | - | - | Soil | Dividices Romulan |
| SRR25508675 | Oak Ridge, Tennessee, USA | 35.98 N, 84.27 W | - | Sediment | Decius Romulan |
| SRR28219342 | Chilgok-gun, Gyeongsang Province, South Korea | 35.96 N, 128.45 E | - | Groundwater | Engmey Klingon |
| SRR6159086 | Lake Superior, Michigan, USA | 47.17 N, 89.66 W | 0.3 | Sediment | Separ + Sud Klingon |
| SRR7028199 | Marcell Experimental Forest, Minnesota, USA | 47.50 N, 93.48 W | - | Peat | Talvath Romulan |
| **Serpens Ridge Vernal Pool Samples (from ggKbase)** |  |  |  |  |  |
| - | Lake County, California, USA (9/9/2021) | 38.69 N, 122.53 W | 1.05 | Wetland | Choi Klingon, Gasko Romulan |
| - | Lake County, California, USA (25/7/2021) | 38.69 N, 122.53 W | 0.9 | Wetland | Chon Klingon |
| - | Lake County, California, USA (25/7/2021) | 38.69 N, 122.53 W | 1 | Wetland | Ghos Klingon |
| - | Lake County, California, USA (9/9/2021) | 38.69 N, 122.53 W | 1.55 | Wetland | Naq Klingon |
| - | Lake County, California, USA (9/9/2021) | 38.69 N, 122.53 W | 1.5 | Wetland | Qel Klingon |

The 45 newly curated ECE genomes were classified into groups using Mauve whole genome alignments, phylogenetics and proteomic presence-absence based analyses **(Supplementary Table 1)**. 23 of these genomes were grouped together and named “Klingons”, respecting the established nomenclature of the Star Trek-based naming convention. Four genomes showed detectable relatedness to Klingons and were grouped together and named “Romulans”. An additional four genomes, which showed moderate similarity to Romulans but were phylogenetically distinct and showed little similarity to anything else in our reference database (ggDB, see methods), were termed “Romulan-like”. The other 14 genomes (11 complete and three near complete) were grouped together and named “Ferengi”.

We identified a single marker protein with detectable similarity across all novel ECEs, and this was used to further investigate the evolutionary relatedness of the three groups. The analysis included many additional contigs that were classified as partial Klingon, Romulan or Ferengi genomes that were not curated in this study, showcasing their breadth. The phylogeny shows that Romulans and Klingons are more closely related than Ferengi are to either **(Supplementary Figure 1).**

### ECEs share gene content with Borgs and their hosts

Of the proteins encoded by these genomes, 42% of Klingon, 18% of Romulan/Romulan-like and 13% of Ferengi proteins had best matches to Borg proteins in the ggDB. Thus, we included sequences from 19 Borgs and 21 related mini-Borgs in our proteome and phylogenetic analyses of the relatedness of all ECE groups **(Supplementary Table 2)**.

In all 42 complete Ferengi, Klingon and Romulan (FKR) genomes we detected perfect terminal direct repeats indicative of circularity, differentiating the FRK genomes from the linear genomes of Borgs and mini-Borgs, which have terminal inverted repeats. The size of the complete FKR genomes ranges from 53 kb to 165 kb, overlapping with the known size range of mini-Borgs (49 - 326 kb, (11)). The Klingon genomes are typically larger in size than those of Romulans and Ferengi, with average lengths of 113 kb (66.4 - 165.5 kb) compared to 76 kb (53.0 - 80.6 kb) and 64 kb (59.5 - 87.0 kb), respectively. The GC contents of all three new ECE families are similar, ranging from 38% - 49%, with an overall average of 44.2% **(Table 2).** This is higher than the average GC of the Borgs or mini-Borgs, which have averages of 32.8% and 33.7% respectively.

**Table 2:**
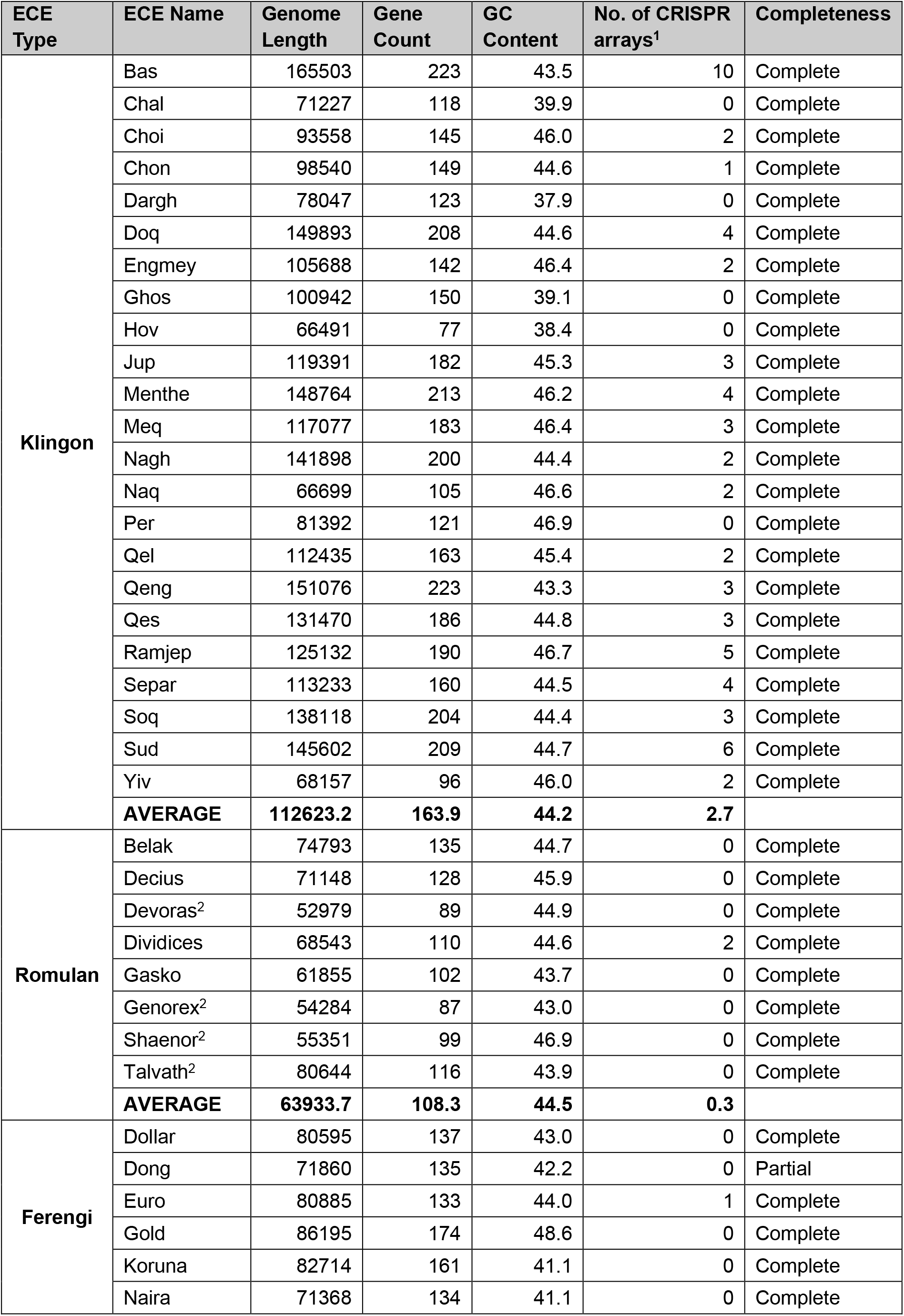
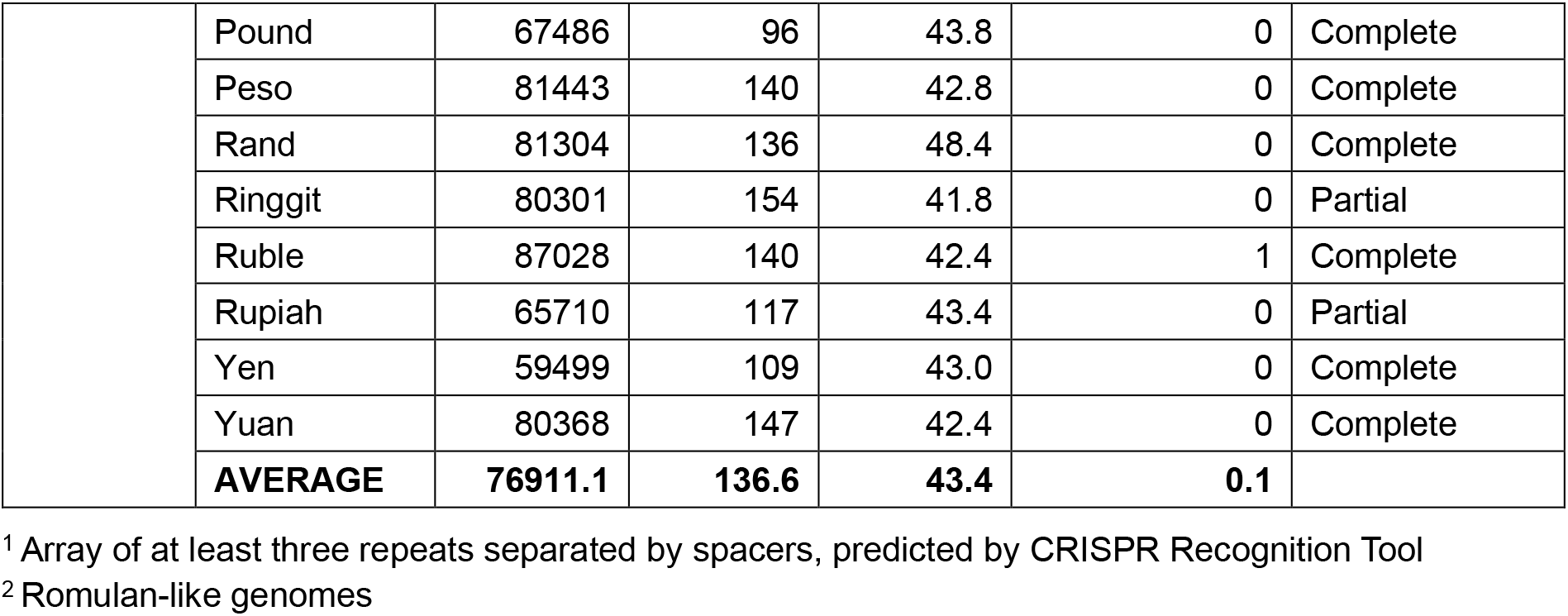
Genomic features of Klingon, Romulan and Ferengi ECE genomes.

To identify the hosts of the novel ECEs, we evaluated the taxonomic profiles of each predicted protein in the FKR genomes and calculated the most common hit. We found Klingons and Romulans shared 14% and 6% of genes with *Candidatus* Methanoperedens, respectively, according to the ggDB database, with an average amino acid identity of 55% and 57% **(Supplementary Table 3)**. Candidatus *Methanoperedens* is the predicted host of the Borg and mini-Borg genomes (8), and within the metagenomes where FKR co-exist with *Methanoperedens*, there are also often the presence of Borg genomes.

CRISPR spacers were also used to link the ECEs to potential hosts. Most of the Klingon genomes contain at least one CRISPR array **(Table 1)**. Eight spacer sequences from the orphan CRISPR-like arrays of five different Klingons matched to *Methanoperedens* genomes **(Supplementary Table 4).** Given all of this, as well as evidence from methylation patterns of hosts and ECEs (see below), we infer that Klingons and Romulans (including Romulan-like elements) replicate in *Methanoperedens* hosts.

Despite their similarity to Klingons and Romulans, Ferengi have very few proteins with best matches or high similarity to *Methanoperedens,* prompting us to look for alternative hosts. The best hits of the Ferengi single copy genes in the NCBI database were consistently from *Candidatus Bathyarchaeota* and Asgard archaea. Protein trees of the resulting matches **(Supplementary Figure 2)** showed the highest similarity to *Candidatus* Bathyarchaeota, leading us to infer that the Ferengi are most likely to be hosted by *Bathyarchaeota*. We specifically looked at the metagenomes that the FKR were found in to pull out four *Methanoperedens* and four *Bathyarchaeota* genomes for inclusion in the proteome comparative and phylogenetic analyses **(Supplementary Table 1, Supplementary Table 2).**

To investigate if ECE relatedness indicates a common ancestor, we used a coalescent approach for phylogenetic inference (see methods; **Figure 2a)**. This enabled us to account for the uneven distribution of genes across ECE types, as only nine protein subfamilies out of 6,197 had a protein representative in all five of the FKR ECEs, Borgs and mini-Borg families, and no single subfamily had a protein representative in every member of two or more ECE types.

**Figure 2:**
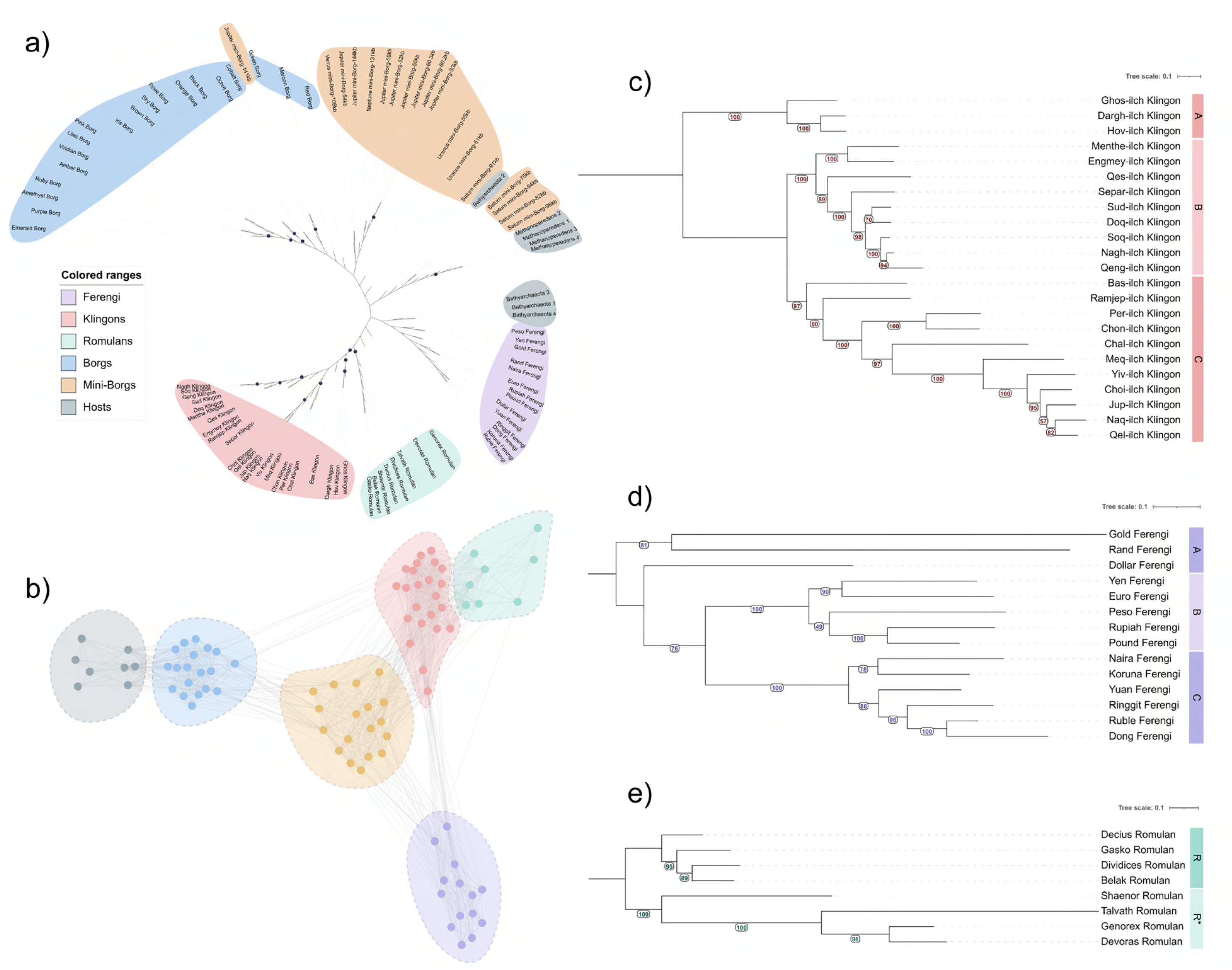
Relatedness of archaeal-associated extrachromosomal elements. a) Coalescence-derived phylogenetic tree displaying the relative relatedness of different archaeal ECE groups. Twelve gene trees were used to derive the coalescent pattern (see methods). Local posterior probability values > 0.60 are marked with navy circles. b) Network of Klingon, Romulan, Ferengi, Borg and Mini-Borg ECEs, as well as predicted hosts, according to shared protein subfamilies. Subfamilies were assigned using HBBLits and MMSeq2. Colour legend is the same as in Figure 2a. c) Concatenated phylogeny of the Klingon ECE family. d) Concatenated phylogeny of the Ferengi ECE family. e) Concatenated phylogeny of the Romulan ECE family. Romulans are marked with R, Romulan-like elements with R*.

All three of the new families formed a monophyletic lineage **(Figure 2a)**. The host genomes included in the phylogeny could not be used to root the tree as they did not form a monophyletic outgroup - most of the *Bathyarchaeota* emerged just before the common ancestor to the Ferengi, suggesting that this common ancestor comes from within the *Bathyarchaeota* genome. However, the *Bathyarchaeota* genomes did not form a monophyletic clade. The *Methanoperedens* genomes were on a clade that also contained Saturn mini-Borg genomes.

The statistical support for the coalescent phylogeny was low, with only 16 out of 90 nodes having a local posterior probability of >0.60. Furthermore, the intra-family topology often did not agree with the concatenated phylogenies built for each ECE family with their shared single copy gene backbone **(Figure 2c-e)**, or in the case of the Borgs, already published phylogenies (9). Phylogenetic analyses using the protein sequences that were inputs for the coalescent were highly discordant **(Supplementary Data 1)**. Based on this discordance, and the near-complete absence of shared genes and therefore lack of a shared genomic backbone, we infer that these five ECE groups do not have a common ancestor. We instead hypothesize that the five ECE families primarily show relatedness as a result of large-scale horizontal gene transfer, rather than vertical descent.

We used amino acid sequences to assign proteins predicted from the genomes of FKR, as well as Borgs, mini-Borgs, *Methanoperedens*, and *Bathyarchaeota*, to subfamilies **(Supplementary Table 1)** and built a protein sharing network based on a presence-absence matrix. Interactions were weighted according to genome size using the Jaccard similarity metric. We bench-marked the maximum Jaccard weighting that showed ECE groups interacting with one another at 0.04. At this value, all groups were interconnected, with Borgs-mini-Borgs, mini-Borgs-Klingons, Klingons-Romulans, and Klingons-Ferengi displaying relatively strong interconnections in the network **(Figure 2b)**. The mini-Borgs have a central position in the network, with strong connections to both the Borgs and all three new ECE families. Above this Jaccard weight, the ECEs began to separate into subsets of interacting groups, with the first interaction to disappear being the one between Klingons and Borgs **(Supplementary Figure 3a)**, and the last interaction to disappear being the one between Ferengi and Klingons **(Supplementary Figure 3c)**.

The arrangement of ECE lineages in the phylogeny as a whole aligns with the arrangement of groups in the protein-sharing network **(Figure 2b)**, supporting the hypothesised strength of each relationship.

The concatenated phylogenies for individual ECE groups generally had high statistical support. In the Klingon phylogeny, 18/20 nodes had confident bootstrap values over 80, and 8/11 in the Ferengi phylogeny **(Figure 2c, 2d)**. In both trees, three monophyletic lineages emerged, labelled clades A, B and C respectively. In the Romulan/Romulan-like element phylogeny, all bootstrap values were above 89, although the Romulan-like elements had extensively long branches despite their shared backbone with the Romulan ECEs **(Figure 2e)**.

Mapping of the genes shared between FKR and Borgs or mini-Borgs revealed a positional bias wherein shared genes tend to be encoded in only one half of the genome **(Supplementary Figure 4)**. This pattern was particularly apparent in the Klingon genomes, although this varied according to clade. In clade A, an average of 26.35% of proteins per genome were assigned to subfamilies that contain at least one Borg or mini-Borg protein, compared with 32.07% of proteins in clade B, and 29.41% of proteins in clade C. In Klingons from Clade B, CRISPR-like arrays and methyl or thiol transferase enzymes were often found at the borders of shared gene blocks.

For Romulans / Romulan-like elements and Ferengi, in over half of the cases, when genes were shared with Borgs, they were also shared with Klingons. The Romulans on average shared more proteins with Borgs (29.08% of proteins per genome) than Romulan-like elements (19.65% genes per genome). This is supported by the protein sharing network, which shows a stronger connection between Borgs-Klingons than both Borgs-Romulans and Borgs-Ferengi, and the almost complete lack of connection between Borgs and Romulan-like elements.

### Replication, transcription and translation of novel ECEs

The GC Skew of each genome was analysed to determine the mode of replication for each ECE. In the cumulative GC skew, some plots show clear troughs and peaks representing the origin of replication and the terminus, respectively **(Supplementary Figure 5a)**, consistent with bi-directional replication. Other GC skew plots were too noisy to make assessments about origin or terminus location **(Supplementary Figure 5b)**. The length of each replichore relative to the genome was variable among ECEs. Unlike the Borgs, each replichore does not encode almost all genes on the same strand.

In detail, the GC skew plots showed local aberrations where the GC skew of one replichore was very similar to that of the other replichore. These regions corresponded almost perfectly with coding strand switches (**Figure 3**). For example, in one Klingon type (Choi), between nucleotides 85,965 and 91,148, the GC skew switches from positive to negative, corresponding to a coding strand switch from forward to reverse strand (**Figure 3a**).

**Figure 3:**
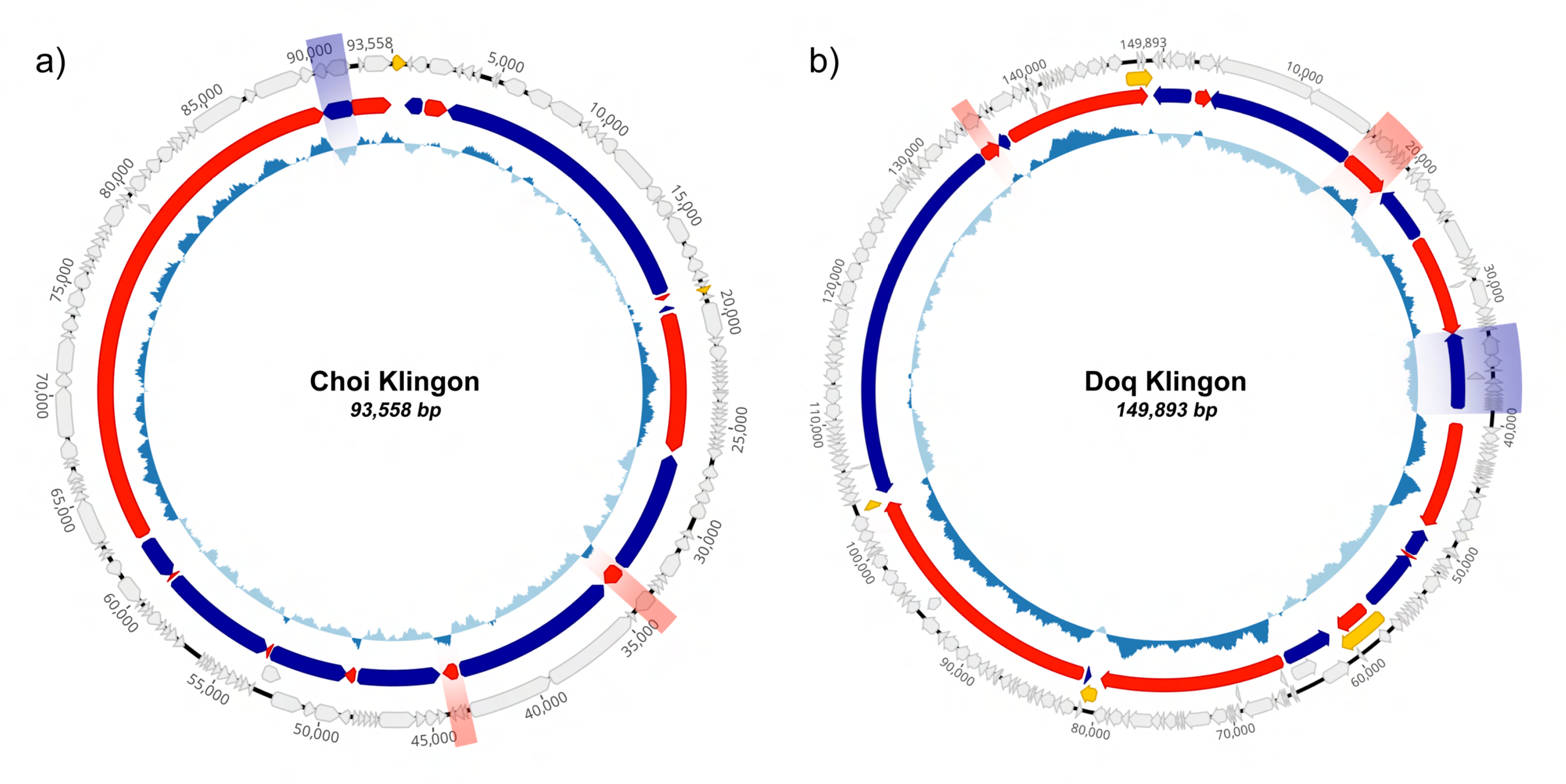
Alignment of gene strand segments with GC Skew plot. Outer-most ring displays genome arrangement, middle ring presents gene directionality (red = forward strand, navy = reverse strand), inner ring displays GC skew (Dark blue = positive skew, light blue = negative skew). Alignment of deviations in strand and GC skew highlighted in dark blue or red of (a) Choi Klingon and (b) Klingon). CRISPR-like arrays marked in yellow.

For Bas, the Klingon with the largest genome, the location of the origin and terminus of replication could not be determined from GC skew due to irregularity in the plot **(Supplementary Figure 5b)**. However, there is clear variation in DNA sequencing coverage across the genome, supporting the inference that the genome replicates bidirectionally. The peak trough ratio (12) and iRep values (13) determined from this plot are 1.55 and 1.59 respectively, indicative of active bidirectional replication at the time of sampling (**Supplementary Figure 5c**).

A paired t-test was performed to analyse whether there was a difference between the average GC content of genes and the GC content of the third codon position (GC3). In Ferengi, there was no statistical difference between the total average GC content vs GC3 (p = 0.9172, t = 0.106). However, the GC3 values were significantly higher than the average GC in Klingons (p = 0.001153, t = 3.7335) and in Romulans/Romulan-like ECEs (p = 0.004757, t = 4.0687). Elevated GC3 was also detected in *Methanoperedens* and *Bathyarchaeota* genomes.

Codon bias and use preference among synonymous codons (relative synonymous codon usage, RSCU) **(Supplementary Table 5)** were calculated for all ECEs and predicted hosts. Cluster 1 in the RSCU matrix contains Klingons, Romulans and *Methanoperedens*, supporting the association between ECEs and host, respectively, and Cluster 2 contains only Ferengi. The *Bathyarchaeota*, two Klingons (Hov and Dargh) and 2 Ferengi (Gold and Rand) were outliers.

A Mann-Whitney U test was performed on the RSCU values to find the most influential codons on the separation of clusters **(Supplementary Table 6)**. The top five most influential codons were calculated to be CGA (Arg, ΔRSCU (C1 - C2) =-0.633), CAA (Gln, ΔRSCU =-0.747), CAG (Gln, ΔRSCU = 0.747), TTG (Leu, ΔRSCU =-0.518) and AGG (Arg ΔRSCU = 1.260). In each case, the Mann-Whitney U statistic was equal to 0 or 396, which indicates complete separation between clusters, and the Benjamini-Hochberg corrected q-value was 2.14 x 10^-6^. Codons that end in a cytosine or guanine are more frequent in Cluster 1 genomes (e.g. CAG, GGG, CCG, CTG and TCC), consistent with the elevated GC3 result, while codons that end in an adenine or thymine are more abundant in Cluster 2 genomes (e.g. CAA, GCT, GTT and ACT).

None of the Klingons, Romulans or Ferengi encode their own ribosomal proteins. An average of 62.6% genes in Klingons encode a Shine-Dalgarno (SD) sequence in their 5’ UTR, followed by 84.4% of genes in Romulans and 49.0% of genes in Ferengi **(Supplementary Table 7)**. Interestingly, all ribosomal binding sites of the clade A Klingons as well as one genome from clade C (Chal) are Bacteroidetes-specific ribosomal binding sites, as opposed to standard SD sequences. In these Klingons, an average of 45.4% of genes contain these binding sites.

### Syntenous single copy genes are unique to each ECE type

We identified protein subfamilies that were statistically enriched in one ECE family relative to the two others to assess what made each family unique. The Klingons have 20 enriched protein subfamilies, over half of which could not be assigned functions. Those functions that could be identified were involved in DNA replication or repair, nucleic acid decoration or were identified as candidates for investigation as viral structural proteins (see below). Romulans have only four statistically enriched protein subfamilies: a recombinase, a clamp loader, and two with unknown functions. In Ferengi there are 16 statistically enriched protein subfamilies, 10 of which have unknown functions. Four enriched Ferengi subfamily proteins contain jelly roll folds, one subfamily is composed of CHAT domain proteins, and one of methyltransferases **(Figure 4a)**.

**Figure 4:**
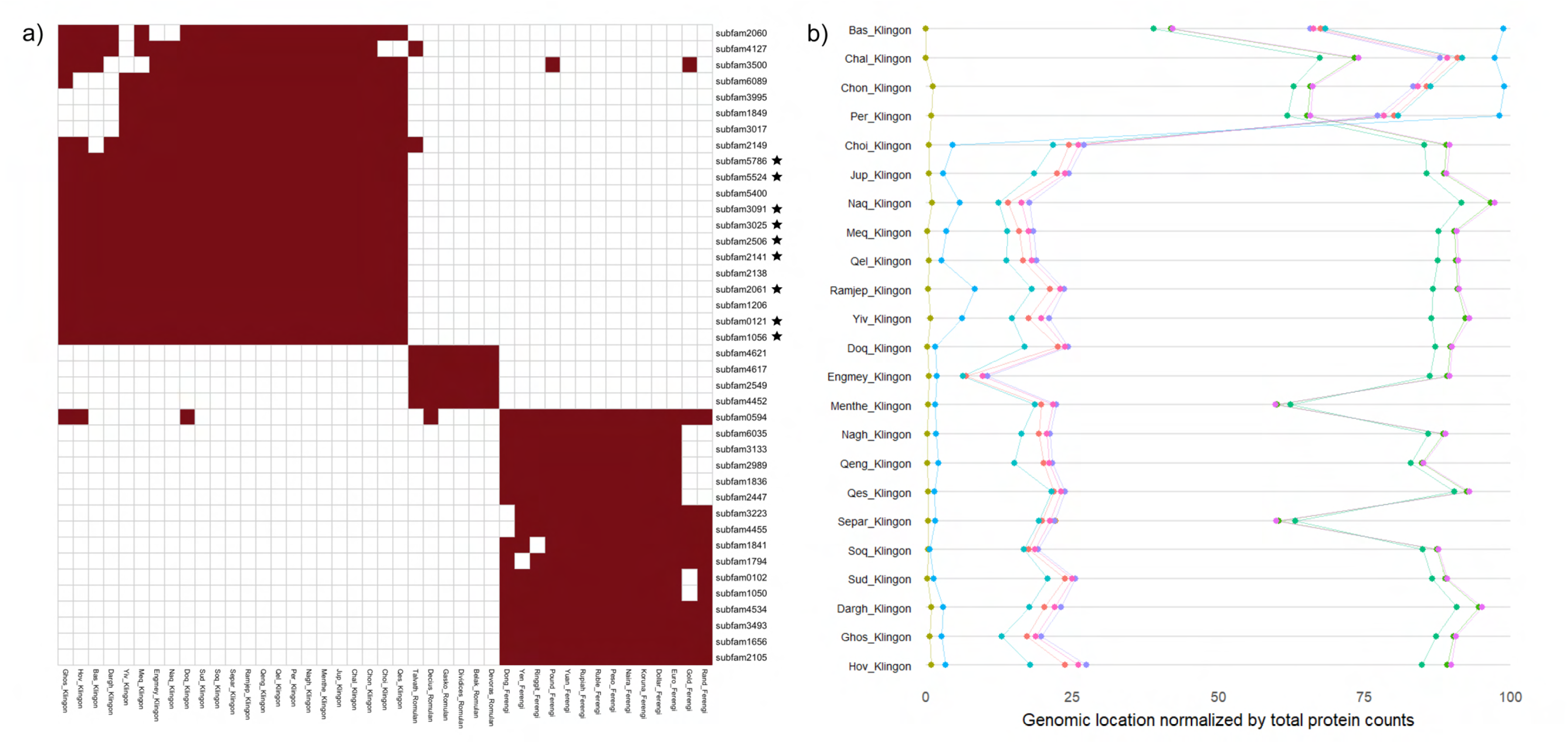
ECE-specific subfamilies and syntenous order. (a) Fisher-test matrix of protein subfamilies for all three ECE families. (b) Relative location of all Klingon SCGs on respective genomes. Klingon SCG subfamilies have been marked in the matrix with black stars.

In the FKR we also identified sets of genes that were present in all genomes of each ECE type and in only one copy, or single copy genes (SCGs). In the Klingons, none of the nine SCG’s subfamilies were shared with proteins from any other ECE type (including Borgs and mini-Borgs), or with their respective putative hosts. All but two of the six SCG subfamilies in Romulans and two of the five subfamilies in Ferengi only contained proteins from that particular ECE type.

We looked at the order conservation of all single copy genes within each ECE type by calculating the Kendall’s Tau (*τ*; (14)) correlation for each ECE pair, and found the SCGs to be almost perfectly syntenous. In Klingons and Ferengi, the median *τ* value was 1. In the Klingons, *τ* was equal to 1 for 140 out of the 153 comparisons; only one large-scale re-arrangement can be seen in four of the Clade C Klingons. (**Figure 4b**). In the Ferengi, conservation of SCG order was perfect in 11 out of 14 Ferengi, with variations in three genomes, two from the divergent clade A (Dollar and Gold) and one from clade C (Yuan) **(Supplementary Figure 6a)**.

The conservation of SCG order was perfect in the Romulans and Romulan-like elements. However, there was variation in the location of SCGs, and therefore the median *τ* was lower (0.33) **(Supplementary Figure 6b)**. However, the presence of a generally shared backbone in all three families supports a model of relatedness via vertical descent from a common ancestor within each ECE group.

### All ECEs have abundant decorative and capsid-like proteins

We folded all predicted protein sequences from Klingons, Ferengi and Romulans using ColabFold and compared each in silico protein model to structural databases via Foldseek to support and extend functional assignments.

To get a better understanding of what protein folds and functions were common beyond matches to PDB structures, we clustered proteins according to their structural similarity. TM-Scores were calculated for 1954 FKR-encoded proteins with confident model predictions (pTM > 0.70; **Supplementary Figure 7**) using an all-against-all structural comparison. 1376 proteins were clustered into 92 groups. The matrix of TM-scores features three clusters of proteins present in ≥ 1 copy in all novel ECE genomes, and six clusters with ≥ 1 representative protein in most (but not all) genomes **(Figure 5, Supplementary Data 2)**.

**Figure 5:**
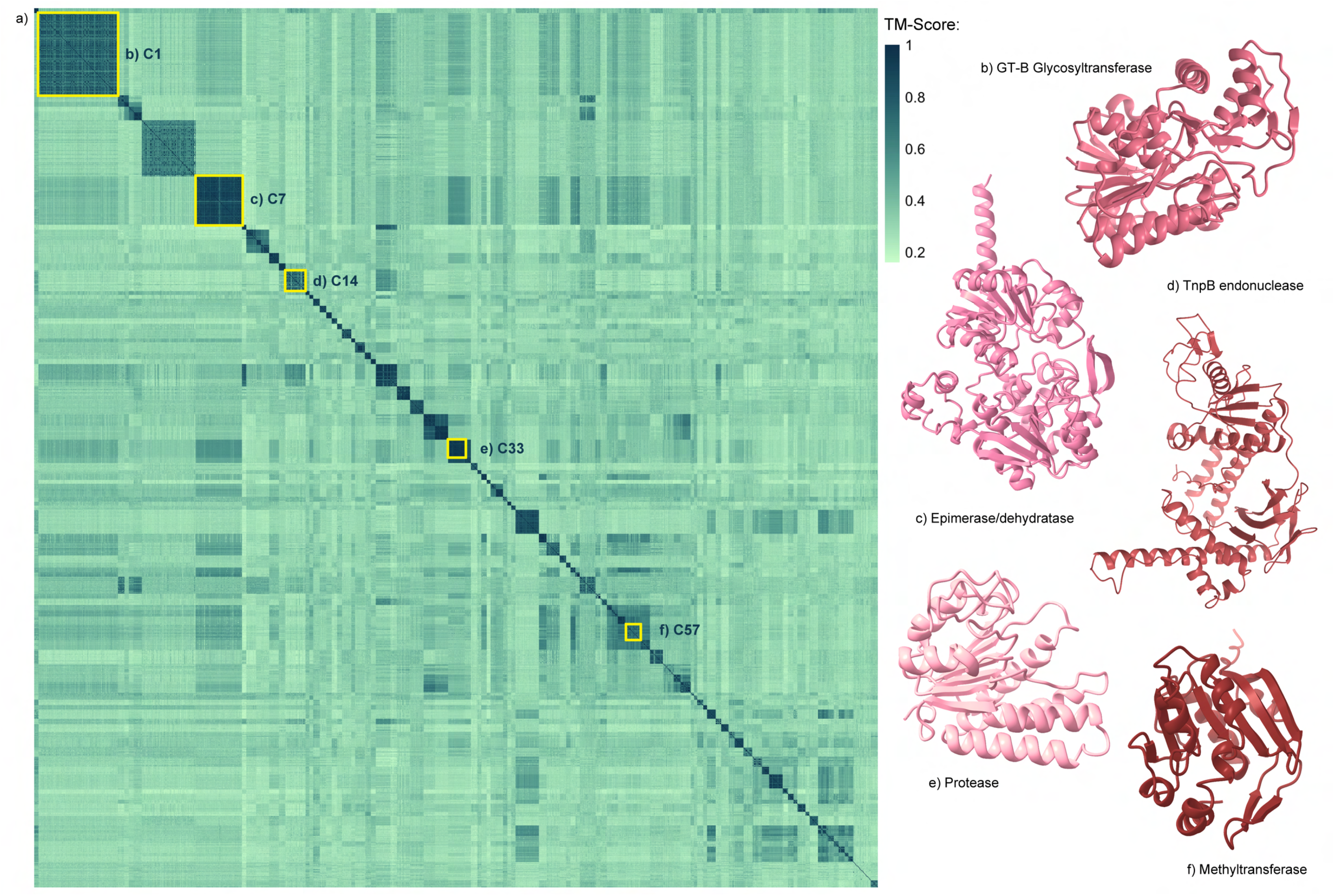
Structural similarity matrix of 1376 Klingon, Romulan, Romulan-like and Ferengi proteins. a) Clustered heatmap showing protein structural similarity according to TM-Score. b-f) Representative proteins of select large clusters. Select large clusters (>= 20 proteins) are highlighted with yellow boxes. Clusters of similar proteins occur on the diagonal. Proteins visualised using ChimeraX.

The three largest clusters, C1, C6 and C7, represent three different types of sugar interacting proteins. The proteins in C1, the largest cluster containing 124 proteins, are predicted to contain two Rossman fold domains joined by a flexible linker and were classified as metal-ion independent glycosyltransferase GT-B type enzymes **(Figure 5c)**.

C6, which has 88 representatives, contains smaller proteins, with only one Rossman fold. These were classified as GT-A type glycosyltransferase proteins that typically depend on metal ions as cofactors and encode a DXD motif for this interaction (15), although this motif was not universally observed.

The proteins of C7, comprising 76 proteins, best match a NAD-dependent epimerase/dehydratase; in particular, GDP-mannose epimerase/dehydratase (GMED; PF16363) **(Figure 5b)**. GMED is an enzyme involved in production of L-Fucose, a carbohydrate that commonly contributes to the membrane profile of the host. Residues involved in NAD interaction, such as the GxxGxxG motif (16) show relatively high conservation across the cluster **(Supplementary Data 3)**.

C33 comprises 29 proteins predicted to be proteinases **(Figure 5e)**. They appear in a single copy in all but one of the Klingons (Chal), and in six of the eight Romulans, with two copies in Decius Romulan. However, there are many more ECE-encoded proteinase proteins that did not fall into this structural cluster as their pTM score was below the threshold for inclusion. The most common PDB match to this cluster (PDB 1CVR; (17)) is arginine-specific cysteine proteinase gingipain R, a protein which cleaves proteins directly after arginine residues. These proteins also structurally align to the CHAT protease domain of SAVED-CHAT filament (PDB 8TL0; (18)), which are known to be activated by target-bound Type III CRISPR effectors, and activate other proteases that initiate dormancy or apoptosis of the host cell (19). This is also the case of C34, which contains an additional nine Ferengi proteins with very strong structural similarity to C33 proteins.

The methyltransferase proteins encoded within Klingons, Romulans and Ferengi did not share enough structural similarity to cluster together in the matrix **(Figure 5f)**. At least five different clusters, C46, C56, C57, C58 and C59 are predicted to contain methyltransferases, with the largest being C57 (25 proteins). There is discernible similarity between them, as indicated by intermediate TM-scores (**Figure 5a** and **Supplementary Table 8**) and by placements in a methyltransferase FoldTree, which also shows different degrees of structural similarity within each cluster (**Supplementary Figure 8**).

Given the abundance of methyltransferases, we attempted to use the DNA modification signal recorded during DNA sequencing to identify potential ECE-host linkages. This was only possible for the Klingon genomes for which PacBio reads were available (Ghos and Choi). For Ghos Klingon only one linkage is indicated, and this is to *Candidatus Methanoperedens* (final score = 0.773), however the shared motifs were relatively short (GATATC and GATC), thus not definitive. Choi Klingon has four matches, three of which are to *Methanoperedens* (final scores = 0.7163, 0.6781 and 0.6766 respectively). In addition to GATC, these linkages relied on RAGTCY in one case and on GGNCCH in the two other cases. Surprisingly, there was one motif match for Choi and an Aminicenates bacterium (final score = 0.8697, shared motifs = GCGATC, RGATCY and SAGTCS) **(Supplementary Table 9)**.

C44 contains proteins from Klingons and Romulans with structural similarity to viral major capsid proteins, suggesting the similarity of these ECEs to archaeal viruses. These proteins correspond to subfamilies 1206, 4620 and 6089. Subfam1206 contains proteins from Klingons, Borgs and mini-Borgs, subfam6089 from Klingons and Borgs, while subfam4620 contains proteins from Romulans only **(Figure 6a)**. Capsid-like proteins often contain jelly roll fold (JRF) domains that are involved in multimerization (20). The order and orientation of the sheets in the Klingon and Romulan proteins varies slightly from the canonical domain structure. They contain nine to ten β-sheets instead of eight, extending out to βK, and they orient themselves into three planes, with five, three and two-three β-strands in each plane, instead of two parallel planes with four β-strands each. Therefore, we refer to these as pseudo-JRF proteins. The two most common PDB matches for these proteins is the VP7/VP4 major capsid proteins from the archaeal viruses HHIV-2 and HCIV-1 (PDB 6H82 and 6H9C; (21)). These PDB structures are also a common best match for Borg capsid-like proteins (10).

**Figure 6:**
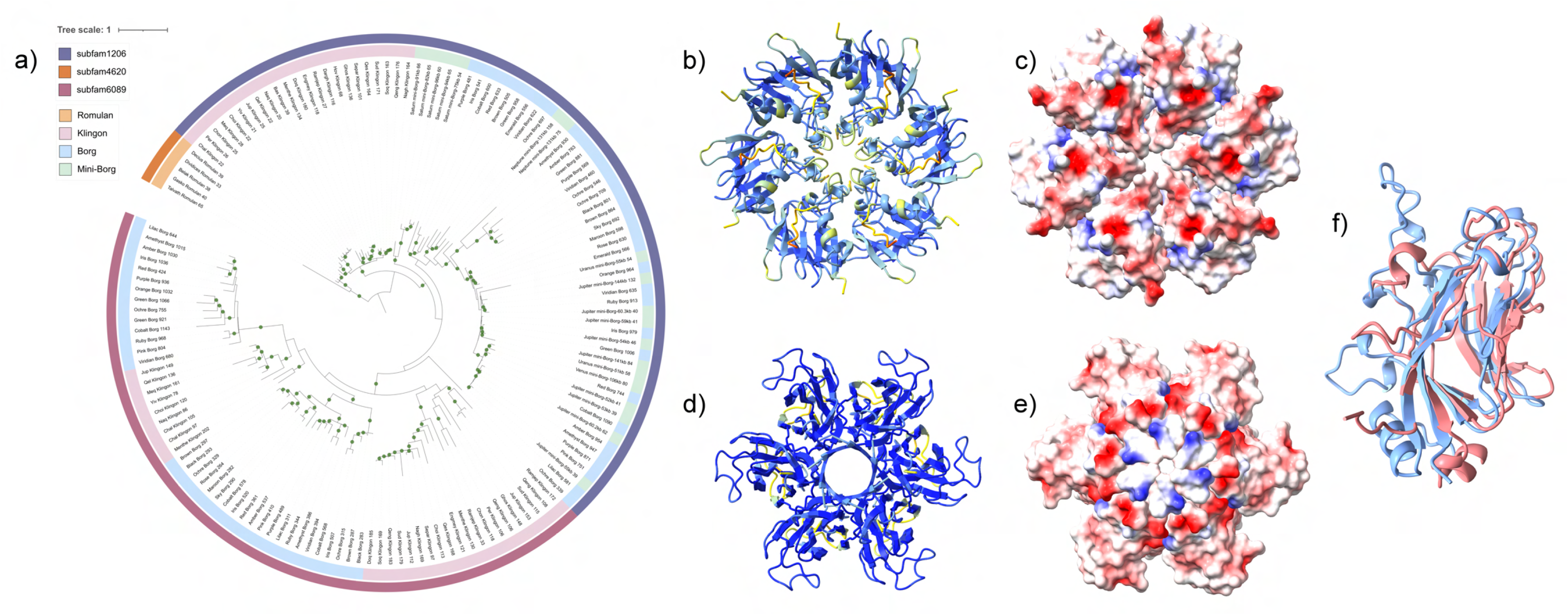
Structure and diversity of capsid-like proteins in Klingons, Romulans and Ferengi. (a) Phylogeny of Cluster 44 proteins, including proteins from subfam1206, subfam4620 and subfam6089. Phylogeny inferred using IQTREE, substitution model Q.PFAM + F + R4. Bootstrap values > 70 are marked with green circles. (b) Homo-hexamer prediction of Chal Klingon gene 22, coloured according to pLDTT (pTM =0.89, ipTM = 0.88). (c) Electrostatic map of Chal 22 homo-hexamer. (d) Homo-hexamer prediction of Ruble Ferengi gene 79, coloured according to pLDTT (pTM =0.79, ipTM = 0.94). (e) Electrostatic map of Ruble 79 homo-hexamer. (f) Structural alignment of Chal 22 with VP7 protein from HCIV-1 (PDB 6H9C).

Ferengi genomes also encode capsid-like proteins, however they do not cluster structurally or at the sequence level with those of Klingons or Romulans. The Ferengi-encoded capsid-like proteins have the expected structure and beta-sheet organisation of canonical JRF domain-containing proteins, and best match the capsid proteins of SPV1 (6OJ0; (22)) and BMV (7PE1; (23)). These JRF proteins correspond with the four sequence-level subfamilies 3223, 4455, 5549 and 3493. Additionally, they align with C75 and C85 of the structural similarity matrix, which collectively contain 32 proteins across 9 Ferengi genomes.

We found that every FKR genome encoded at least one protein with a JRF or pseudo-JRF domain that was annotated as a capsid protein, although not all met the pTM inclusion threshold for this analysis.

The HCIV-1 and HHIV-2 capsid proteins naturally form hetero-hexamers (24). We tested for multimerization the capsid-like proteins identified in the subfamilies listed above, although this does not include every capsid-like protein encoded by FKR. Of the 21 proteins from subfam1206 and three from subfam4620, all but two had more confident ipTM scores as hexamers compared to pentamers. All proteins multimerized as hexamers with ipTM scores of > 0.6, with 18 proteins having scores that ranged from 0.76 to 0.88 **(Supplementary Table 10)**, with the most confident being the model for the homo-hexamer for Chal Klingon gene 22 **(Figure 6b)**. One face of the homo hexamer has a clear negative charge **(Figure 6c)**. Despite adopting pseudo-jelly roll folds and clustering with C44, there is no strong evidence for multimerization of the 14 proteins from subfam6089. One protein’s pentamer prediction (Per Klingon gene 106) had an ipTM of 0.61, but all others were < 0.5. In Ferengi, proteins from subfam5549 had more confident pentamer predictions, and all others had more confident hexamer predictions, reaching ipTM values of up to 0.94 **(Figure 6d)**. These multimers also are predicted to have a negatively charged face once assembled **(Figure 6e)**.

Given that all of the FKR genomes encoded at least one capsid-like protein, we suspected that the ECEs may be viral. This classification was supported by GeNomad metrics **(Supplementary Table 7)**. However, we were unable to identify any DNA packaging machinery, such as terminases, in most of the FKR genomes. The one exception is AAA-domain containing ATPases, which were identified in the three clade A Klingons and may be involved in packaging, however this was far from universal.

Common structural annotations from the Foldseek screen that did not cluster together in the matrix include antitoxins from toxin-antitoxin systems (e.g. 5WZ4 (25), 7YCW (26)), DNA repair proteins (e.g. 2QLC, (27)), ubiquitin ligase (e.g. 2RJ2 (28), 6H77 (29)) and XerA recombinases (e.g. 5HXY, (30)).

Altogether, FKR genomes share the capacity for genome or membrane decoration with methyl and sugar compounds. They all contain proteins related to programmed cell death and encapsulation (consistent with extracellular transfer), suggesting that they do not permanently reside in single host cells.

### Orphan CRISPR arrays and novel TnpBs

Most Klingons encode orphan CRISPR-like arrays (i.e., lacking canonical Cas effector proteins), which are rare in Romulans and Ferengi **(Table 2)**. 17 of the 23 Klingons, as well as two Ferengi (Euro and Ruble) and one Romulan (Dividices) contain CRISPR arrays with ∼30 bp repeats separated by unique spacers, also ∼30 bp in length. The number of arrays with at least three repeats, and number of spacers per array, vary among genomes, with up to 10 arrays (Bas Klingon) and up to 144 spacers over six CRISPR regions (Sud Klingon). These arrays are often, though not always, positioned in between gene strand switches and often align with aberrations in the GC skew expected for a bidirectionally replicating genome **(Figure 3b)**.

As Cas proteins were not identified in any of the ECE genomes, we suspected that the ECE systems rely on host machinery to acquire and make use of CRISPR spacers. The *Methanoperedens* hosts encoded Cas1 and Cas2 proteins, but these proteins were not detectable in the *Bathyarchaeota* genomes. As use of host machinery likely depends on similar repeat sequences (31, 32), we searched for matches between orphan CRISPR and host array repeats, but no matches were identified. Additionally, we were unable to detect any similarity at the secondary structure level between repeats of FKRs and of *Methanoperedens* hosts.

Four Klingon arrays (one from Bas, one from Ramjep and two from Yiv) contained repeats that could be classified according to their CRISPR system type (see methods). Those from the Yiv Klingon have best matches to a Type V-K, the array from Ramjep to a Type I-A and the array from Bas to a Type I-B system. The only category of CRISPR array that could be resolved from the *Methanoperedens*’ genomes with over 70% probability was Type I-D systems in two of the eleven arrays tested.

It was already known that spacers in the *Methanoperedens* CRISPR arrays map to Klingons - specifically, two spacers which map to the middle of a transpose gene in Choi Klingon (5). We found that this relationship is reciprocal, as some of the Klingon-encoded spacers also match *Methanoperedens* genomes **(Supplementary Table 4)**, one of which belonged to an array that was classified as a Type V-K. In most cases, the spacers target within a gene. Interestingly, where functions of the encoded proteins could be identified, many of the functions are essential or important for defence, e.g., cytochrome, DNA helicase, restriction endonuclease. The sequences targeted by the spacers were more commonly found closer to the 5’ end of the respective gene than the 3’ end. No candidate omega RNAs (as identified in the Borgs; (10)) could be discerned in any FKR genome. We were unable to identify any additional *Methanoperedens* spacers that mapped to a FKR genome, although interestingly we did identify a match to an Aminicenanates bacterium, the same phylum as was identified in a potential host of the Choi Klingon in the methylation host-ECE analysis (e-value = 6.00e-06).

Five of the Klingon genomes encode proteins and nearby arrays of the new VIPR defence system reported by Yoon et al. (33). None of the predicted VIPR RNAs, predicted using the rules of Yoon et al., target any sequence in any metagenome from the sample site.

We identified putative TnpB-like proteins in all but four of the FKR genomes containing orphan CRISPR arrays, some but not all of which were encoded adjacent to an orphan array. Consistent with this identification, *in silico* protein structure models show a strongly positively charged transposase (PF01385) domain, all contain a zinc finger-containing DNA binding domain, and most of the enzymes contain a conserved DED/DDE box within a RuvC domain that is essential for DNA cleavage **(Supplementary Figure 9, Supplementary Data 4).**

A subset of these TnpB-like proteins comprises C14 in the structural similarity matrix **(Figure 5d)**. C14 is composed of 33 proteins, of which 31 were assigned to eight sequence-level subfamilies (two from Ferengi were unique and not assigned to any subfamily). All except one of these eight subfamilies contain TnpB sequences from Borgs. The single Klingon specific subfamily is subfam1239 and contains 16 proteins. These proteins contain DNA binding domains but do not encode the conserved DED residues in the same position of the RuvC domain compared to the other TnpB proteins. Instead, there is typically a conserved glutamate or aspartate residue a few (∼3-5) amino acid positions away **(Supplementary Data 4)**, but these do not come together in three-dimensional space with the other D/E residues. This suggests that these TnpB-like proteins can bind to DNA but not cleave it.

While many of the TnpB proteins have significant hits to other transposases in the NCBI dataset, some have no matches with percent identity > 70%, and most only have matches to sequences from their host genus (either *Methanoperedens* or *Bathyarchaeota*), indicating that these proteins are relatively unique to their host’s genome. Their abundance in FKR genomes and frequent co-occurrence with orphan CRISPR arrays suggest involvement in host-ECE and possibly even ECE-ECE interactions.

## Discussion

In this study, we report 45 novel ECEs and assign them to three new families, Klingons, Romulans and Ferengi, expanding the known diversity of archaeal ECE types. These ECEs are very prevalent in some ecosystems, suggesting substantial potential to impact the activity and/or fitness of their hosts. The Ferengi were identified from groundwater samples, whereas Klingons and Romulans were found in many different types of environments. The circular genomes of Klingons, Romulans and Ferengi reach sizes larger than the largest archaeal viruses and help bridge the gap between canonical archaeal plasmids and viruses and the giant linear Borgs (8). Phylogenetic analysis, CRISPR-spacer matches and analysis of shared protein subfamilies indicate that the Klingons and Romulans are hosted by *Candidatus Methanoperedens*, whereas protein similarity analyses suggest that the Ferengi are hosted by *Bathyarchaeota*. The assignment of *Methanoperedens* in the case of Klingons and Romulans is also supported by GC/GC3 and codon usage statistics. We hypothesize the FKR genomes replicate bi-directionally, similar to other dsDNA circular phage and eukaryotic viruses (34).

From the phylogenetic analyses, we infer that the apparent relatedness within FKR and between them and the Borgs stems primarily from horizontal gene transfer, rather than vertical descent from a common ancestor. The structure of the protein-sharing network suggests that the relation between all five ECE groups is indirect, with mini-Borgs acting as a central source of shared genes that are horizontally transferred to Borgs, Ferengi and Klingons - the latter of which are then passed onto Romulans.

The aberrations in GC-skew that align with coding strand switches within the genome may be indicative of large-scale inversion mutations that would flip the GC signature (35). Alternatively, these blocks of genes could be remnants of horizontal gene transfer. Frequent and recent horizontal gene transfer between FKR ECEs and Borgs / mini-Borgs would explain their shared genetic content despite the lack of a similar backbone. In any case, these new ECEs appear to be key vectors for horizontal gene transfer amongst the archaeal communities, and large scale mutations appear to be common.

In some of the ECEs we identified potentially novel CRISPR-like systems that combine TnpB genes and arrays. CRISPR elements are frequently horizontally transferred among organisms (36), and thus the FKR arrays may have been acquired by horizontal gene transfer along with nearby genes. The presence of (often several) CRISPR arrays lacking Cas machinery is a notable feature of the Klingons and some Romulans and Ferengi, and we speculate that if the CRISPR-arrays are active in FKR genomes, that they are using the FKR-encoded TnpB proteins as effectors, rather than host-encoded machinery. CRISPR arrays are known in ECEs, such as large phage, where they typically have an effector protein or proteins but lack acquisition machinery (37). In these cases, it is likely that addition of spacers to the phage loci involves host Cas1 and Cas2 enzymes, requiring that the host and ECE CRISPR repeats are compatible. In the current study there was no detectable similarity between host and ECE repeat sequences in their sequence or in the repeat secondary structure. Thus, the means by which spacers are acquired into the archaeal ECE arrays remains unclear. Greater confidence about exact host-ECE pairings may help better explain the origin of these arrays.

Despite the fact that FKR genomes lack replication, transcription or translation machinery and likely rely on their hosts for propagation, the presence of spacers targeting *Methanoperedens* in select CRISPR arrays from the Klingons suggests host antagonism. One explanation is that the ECE-encoded TnpBs, some of which are nuclease dead, are using the CRISPR spacer sequences to silence the expression of host metabolic, replication and defence-related genes. Targeting the 5’ end of the gene could block the binding of RNA polymerase, preventing expression of anti-viral defence genes. Transcriptional silencing is associated with Type IV CRISPR systems, which also lack the Cas1 and Cas2 acquisition proteins (38). Alternatively, the TnpBs and spacers may be used by ECEs for abortive immunity upon co-infection with competitor ECEs. Finally, the TnpBs may both silence the expression of replication related genes to induce dormancy of their host and suppress replication of competitor ECEs. Initiation of host dormancy has been observed in multiple systems, such as PemIK in *Klebsiella* and the ToxIN system (39), and would evolutionarily be more favourable than total cell suicide. Testing the enzymatic activity of these spacer-guided TnpBs, both those with full DED boxes involved in nuclease activity and without, may clarify whether they cleave DNA and are functioning primarily as nucleases or transcriptional regulators.

The presence of addictive toxin-antitoxin systems in Klingons, Romulans and Ferengi suggest that they are selfish-acting elements that establish long-term associations with their hosts. Consistent with this is the elevated GC content of the third codon position (GC3) in Klingons and Romulans, which is known to indicate long-term host association (40). As the third codon is most flexible to mutation, elevated GC3 may develop following the introduction of a new ECE into a new host. Alternatively, this may be important in cases where regions of the ECE genome have lower GC content than that of the host. As many of the genes of the new ECEs, especially Klingons and Romulans which were noted as preferring codons that end in guanine or cytosine, appear to have been horizontally transferred from Borgs (which have considerably lower GC than *Methanoperedens*), the elevated GC3 may be the consequence of gene domestication.

Capsid-like proteins are the only protein universally encoded in FKR, Borg and mini-Borg genomes, indicating they are key to their existence. The capsid-like proteins may be involved in encapsulation, potentially for environmental persistence external to a cell, and/or transfer between hosts. As with Borgs (10) and mini-Borgs (5), the lack of universal packaging machinery prevents their classification as canonical archaeal viruses. The FKR and Borgs are not the only archaeal ECEs to encode capsid proteins but no terminases. Modeling of seven predicted structural proteins enabled a full virus reconstruction for ∼4 - 5 kb Charybdis elements of *Poseidoniales* archaea from marine environments (41). The authors reported that these uncultivated viruses also lack typical virus hallmark proteins, such as terminases. Given that the Charybdis elements, Borgs and mini-Borgs, and now FKR elements, all share this pattern, we hypothesize that the method for packaging DNA in these archaeal viruses is independent of terminases. For example, the capsid may assemble around the viral genome, similar to nuclear-forming jumbo phage such as ΦKZ (42) and then exit the host cell via budding or a similar mechanism.

The other uniting feature of the FKRs is the abundance of genes involved in decoration, such as glycosyltransferases, putative GDP-mannose epimerase/dehydratase (GMED) proteins and methyltransferases. These enzymes may be used to modify the ECE genomes or decorate their capsids. Alternatively, a subset may modulate the host epigenome or alter the membrane profile of their hosts. Suggesting a role in self-modification, the PacBio data indicate that Klingon DNA contains some form of epigenetic modification, possibly methylation, although it is unclear whether this is generated by ECE-encoded or host-encoded methyltransferses.

The presence of GMED implicated in fucosylation is curious, as fucosylation of archaeal cell membranes is exceedingly rare (43). Given this, and the diversity of sugar related enzymes encoded in ECE genomes, we hypothesize that FKRs produce capsids decorated with molecules also found on host cell surfaces. Resembling the host cell surface may be important for host identification and infection. If the FKR are enveloped viruses, diverse surface glycopolymers may enable identification of diverse hosts, and may imply a broad host range.

Evidence for many horizontally transferred genes suggests that the FKR are important vectors for transfer of genes between themselves, other ECEs and their hosts, thus can contribute to both microbial and ECE evolution. An important conclusion of this study is that gene transfer is so prevalent that it creates the impression of relatedness between these elements that is not supported by phylogenetics. The transfer of genes among ECE types likely implies co-occurrence in a host cell, at least for part of their life.

## Conclusion

Although many ECEs genomes can now be reconstructed from metagenomic datasets, they are often unlinked to host populations. Thus, there is generally scant knowledge of the suite of ECEs that associate with any specific group of hosts, particularly for hosts that are not available in pure cultures. This is especially true for archaea. Klingons and Romulans expand the previously described group of *Methanoperedens* ECEs, composed of Borgs, mini-Borgs, plasmids, viruses, mini-chromosome-like elements (1, 5, 8, 11). This makes *Methanoperedens* an interesting case study for understanding of the nature of archaea - ECE associations. The introduction of Ferengi also unveils that the study of archaeal ECEs may go beyond what can be observed with a single host organism, and elucidates the true degree of ECE-enabled gene transfer that influences the evolution of archaeal microbial communities.

## Methods

### DNA Sampling, Sequencing and Processing

Soil samples were collected from a wetland site in Lake County, California, at depths of 90 - 155 cm. DNA was extracted using Qiagen DNeasy PowerMax Soil Kit and sequenced using Illumina and PacBio technologies. Raw short reads were processed using BBDuk (44) to trim or remove low-quality sequences, then assembled using metaSPAdes v4.2.0 (45). HiFi long reads were assembled using metaMDBG v0.3 (46).

### Public Database Search

Given that the Klingon host is inferred to be *Methanoperedens* (5), we searched for the presence of Klingons and related ECEs in the public datasets where *Methanoperedens* was detected. In brief, Sandpiper (47) was leveraged to identify metagenome samples containing *Methanoperedens*, whose assemblies were obtained from Logan (48). For samples absent in Logan, raw sequencing reads were downloaded and classified using GraftM (49) with a custom package (11). Samples that contain Klingons, Romulans, Romulan-like elements of Ferengi were processed and assembled following the protocol above.

Open reading frames of assembled scaffolds were called by Prodigal v2.6.3 (50) using genetic code 11. Scaffolds that encode FKR marker genes and proteins shared with previously reported genomes (5) were considered as potential FKR ECEs.

### Assembly and Curation of genomes

The potential Klingon and related ECE genomes were curated using Illumina and PacBio reads, as detailed in Shi et al. (11). Briefly, reads were mapped to scaffolds with 5% mismatches to check and fix assembly errors. The reads placed at scaffold ends and their unplaced paired reads (in the Illumina sequencing) were used to extend the sequences. The extended scaffolds sometimes merged with other scaffolds based on overlap. In some cases, reads extending one scaffold side matched back to the other side, indicative of circularization of scaffolds. Reads were then mapped with 0% error to check if the entire scaffold was covered by reads evenly.

The start of each curated genome (i.e. where gene numbering started) was set based on the presence of conserved syntenous gene blocks identified using Mauve whole genome alignments. In cases where GC skew plots were readable, these were taken into consideration and Mauve blocks closest to the estimated origin were selected as the start point.

### Annotation

Functional annotations for open reading frames identified by Prodigal were assigned through comparison with the Kegg (51), UniRef (52) and UniProt (53) databases by DIAMOND v2.2.5 (54). Proteins discussed in detail in this manuscript were also compared to the NCBI CDD v3.21 (55) and Pfam v37.2 (via HMMER; (56)) databases for domain annotation. To identify the closest taxonomic match of proteins, they were compared to the ggDB database, composed of sequences and taxonomic assignments from the Kegg, UniRef and UniProt databases. This database was also used to calculate the best taxonomic match for each gene encoded in the FKR **(Supplementary Table 3)**.

### Measuring ECE Abundance within metagenomes

The 50 highest coverage values were extracted from each metagenome and compared to the coverage of each curated ECE genome. Coverage values were calculated using Bowtie2 v2.5.4 (57). Coverage of the ribosomal protein S3 was used for each contig. In cases where there was more than one ECE per metagenome, ECE coverages were summed. ggDB was used to identify the domain taxonomy of the top five non-ECE contigs per metagenome.

### Protein sequence clustering

Amino acid sequences from all Klingon, Romulan and Ferengi genomes, along with those from Borgs, mini-Borgs, four *Methanoperedens* and four *Bathyarchaeota* genomes were extracted. These were clustered into subfamilies according to amino acid sequence using HHBlits v3.1.0 (58) and MMSeqs2 (59) based on a method described in Méheust et al. (60) **(Supplementary Table 1)**.

The subfamily classifications were used to create the ECE-interaction network. A Jaccard Index (61) was applied to account for differences in genome size. Jaccard weightings between 0-0.2, in intervals of 0.01, to investigate the strength of connectivity between ECE groups. The network was visualised using the ggplot2 (62), ggraph (63) and igraph (64) packages on R (65) via RStudio v2026.01.0 (66).

Fisher’s exact test was performed to identify enriched subfamilies per ECE type. The resulting matrix was calculated and visualised using the tidyverse (67) and ggplot2 R packages.

### Phylogenetics

To produce the coalescence-based phylogeny, protein subfamilies were ranked according to their abundance across Klingon, Romulan, Ferengi, Borg and mini-Borg genomes. We set four strict criteria to select subfamilies to build into protein trees, to use as input for the coalescent phylogeny. The subfamilies had to contain proteins from at least three different ECE families, had to be present in at least 10 genomes total, 65% of the proteins had to exist in a single copy per genome and the maximum copy number allowed was four. A swap-based local search was then performed to select these subfamilies, maximising mean taxon occupancy, evenness of mean occupancy and minimising cases of multiple copies per genome. This resulted in the selection of twelve subfamilies: subfamilies 0594, 1206, 1444, 1731, 1781, 2036, 2063, 2448, 3931, 5400, 5532 and 5921 **(Supplementary Table 1)**.

Subfamilies were aligned with MAFFT-linsi v7.505 (68) and trimmed with TrimAL v1.5 (69) The single-protein phylogenies were built using IQTREE’s Model Finder Plus (70, 71) algorithm. Substitution models were selected according to BIC scores. The models chosen for each subfamily were, respectively, LG+I+G4, Q.YEAST+F+I+R4, LG+F+R5, LG+F+I+R5, LG+I+G4, LG+G4, LG+R4, LG+I+G4, LG+I+G4, WAG+F+G4, LG+I+G4 and LG+G4.

ASTRAL-Pro3 from ASTER v1.25 (72) was then used to infer the coalescent phylogeny and predict local posterior probabilities for each node, due to its ability to account for paralog genes in each protein tree. 16 rounds of search and subsampling were performed for phylogeny inference. Due to difficulties in identifying the root position in gene trees, due to the non-monophyletic nature of host-encoded “outgroup” proteins, we chose to portray the resulting phylogeny as unrooted. Furthermore, as ASTRAL-Pro3 requires accurate rooting for determination of branch lengths, we chose to ignore branch lengths in the figure.

For the ECE-specific phylogenies, a concatenated approach was used. IQTREE was used in each case, with 1000 ultrafast bootstraps. The substitution models used for the Klingon, Ferengi and Romulan-only trees were Q.INSECT+F+R4, Q.PFAM+F+I+G4 and Q.PFAM+F+G4 respectively. This method was also used for the inference of the capsid-like protein phylogeny, (Q.PFAM+F+R4 substitution model), the Ferengi-host phylogenies (VT+F4, WAG+G4 x 2 and Q.YEAST+G4) and the partial FKR scaffold tree (LG+I+R5 substitution model). In all cases, IQTREE’s ModelFinder was used to determine the most appropriate substitution model.

For the Ferengi-host phylogenies, a representative sequence with the highest average pairwise identify to all other proteins in the subfamily, from each Ferengi SCG subfamily, was chosen and input into BlastP (73). All significant hits were extracted, and phylogenies were inferred for each subfamily. Hits in the smallest possible monophyletic clade containing all Ferengi were inspected manually to assess their taxonomy.

FoldTree (74) was used for inference of the methyltransferase structure phylogeny.

Determination of the root position of rooted phylogenies displayed in this manuscript were chosen according to pairwise similarity of genomes, as measured by number and size of blocks generated by Mauve whole genome alignments, or by manual inspection of single-protein alignments. All phylogenies were annotated and visualised via iTOL v7.6 (75).

### Genome Characterisation and Arrangement

The inspection of genome architecture and arrangement was done within Geneious Prime v2025.1.3 (76). To identify shared genes between Borgs / mini-Borgs and ECEs, subfamilies with at least three Borg representative proteins and three proteins from the ECE in question were extracted. The proteins present in these subfamilies were then manually annotated on circular genome maps in Geneious.

To calculate GC skew metrics, the iRep (13) package was used. GC Skew plots were generated in Proksee (77). To identify the type of ribosomal binding site, GeNomad v1.12.0 was used (78). GeNomad was also used to predict the nature of ECEs as either chromosomes, plasmids or viruses.

The Biopython package (79) was used to calculate CG3 content, codon bias and RSCU values for every annotated gene. The RSCU matrix was designed to look at all sense codons. HDBScan (80) was used to cluster genomes according to their RSCU, and the wilcox.test function in R was used to generate statistics for the Mann-Whitney U statistical test to identify codons that contributed most to the HDBScan-generated clusters.

Subfamilies with exactly one protein per ECE per ECE family were extracted and classified as SCGs. The location of these proteins was recorded and a Kendall’s *τ* (81) statistical analysis was performed to calculate whether the relative location of the SCGs was statistically similar to other ECE genomes. Once statistical significance was confirmed, the proteins were mapped on linear genome maps with the dplyr (82) and ggplot2 R packages. The start position of each genome was modified for the purpose of figure creation so that SCGs were positioned on the same side of the start point (to account for the circular genomes being represented as linear, which would otherwise position SCGs on opposite ends).

### Protein modelling and clustering

Protein structures from all Klingons, and Romulans, and select Romulan-like and Ferengi genomes were predicted via local runs of Colabfold using parameters ‘—amber --templates --num-recycle 3’ (83). Only proteins from select Ferengi were folded based on the genomes that had been curated at the time of analysis. To assess convergence of the five protein models generated by Colabfold for each protein, pTM scores for each model were extracted and a rainfall plot was generated and visualised in R using ggplot2 and dplyr packages **(Supplementary Figure 5)**.

Protein models with pTM > 0.70 were extracted for structural clustering analysis. TM-align was used to generate TM-Scores for all possible protein comparisons. This was arranged into a matrix which was clustered by the python HDBScan package.

For functional annotation, these models were initially compared to proteins in the Protein Database (84) via FoldSeek (85). Only functional annotations where e-value < 0.05 was considered. Protein structure and alignment with Foldseek matches were also manually inspected for each protein discussed in detail in this manuscript using UCSF ChimeraX v1.10.1 (86).

Multimer models for the capsid-like proteins were generated with Alphafold3 using default settings (87). Multimers of 2-12 subunits were analysed to assess the most confident configuration of monomers.

To search for conservation of residues within clusters discussed in this manuscript and to visualise alignments, we used Jalview v2.11.5.2 (88). To identify conserved DED residues in TnpB enzymes, they were compared to known TnpB references from (89) and (90).

### Methylation analysis

For the two metagenome datasets where PacBio datasets were available, modification analysis was performed using MODIFI (91) to predict modified motifs and predict host-ECE linkage. Only results with a final score > 0.60 and specificity < 0.001 were considered.

### Identification and analysis of repeat arrays

Putative CRISPR and VIPR arrays were identified using the CRISPR Recognition Tool (CRT) Geneious Plugin (92) with a minimum repeat number of three. The CRISPR arrays were then further validated with the PADLOC web server (93). Spacer sequences from these arrays were extracted using the BEDTools command line software (94) and matches from within the respective metagenomes were searched using blastn-short. Classification of CRISPR systems according to their repeat sequences was performed using CRISPRCasTyper (95).

## Declarations

### Ethics approval and consent to participate

Not applicable

### Consent for Publication

Not applicable

### Availability of Data and Materials

FKR sequences are publicly available via https://ggkbase.berkeley.edu/project_groups/fkr_genomes. All Supplementary Data are available via https://doi.org/10.5281/zenodo.23050150. All scripts used in this study are publicly available via https://github.com/njablonsky/FKR_Project.

### Competing Interests

All authors declare no competing interests related to this work.

### Funding

This research was supported by an Australian Government Research Training Program (RTP) Scholarship at Monash University to NJJ, the Innovative Genomics Institute at Berkeley and the US Department of Energy via Lawrence Berkeley National Laboratory to JFB, and the Snow Medical Research Foundation (SMRF2021-276) to GJK. None of the funders were involved in the design of the study, data collection, analysis, interpretation of the data, the writing or the decision to submit the article for publication.

### Author Contributions

This study was designed by NJJ, JFB and GJK. Searching of public data for genomes and genome curation was performed by L-DS and JFB. All metagenomic composition, phylogenetic, protein folding, gene content, functional annotation, protein structure clustering, GC skew and genome arrangement analyses were performed by NJJ with input from JFB. Protein sequence clustering, single copy gene order and synteny analyses were performed by NJJ, with input from L-DS. Extraction of partial ECE scaffolds was done by L-DS. PacBio DNA Modification analysis was performed by SW, with input from RS. TnpB analysis was performed by NJJ, with input from DAG, LVA and GJK. VIPR analyses were performed by NJJ with input from LVA and JFB. The manuscript was written by NJJ and JFB, with input from all co-authors.

## Supporting information

Supplementary Figures 1 - 9 and Summary of Supplementary Information

## Acknowledgements

We thank Professor Fasseli Coulibaly, Dr. Marjan Hadian-Jazi, Loki Sangalli, Kinsey Long, Susan Mullen and Leylen Miloslavich for their helpful input. We also thank all members of the Banfield and Knott labs for their useful suggestions.

## Notes

https://github.com/njablonsky/FKR_Project

https://ggkbase.berkeley.edu/project_groups/fkr_genomes

https://zenodo.org/records/23050150

### Competing Interest Statement

The authors have declared no competing interest.

