## Supplementary Figures 1 - 9 and Summary of Supplementary Information for "Genetic exchange drives relatedness of large linear and circular extrachromosomal elements of archaea"

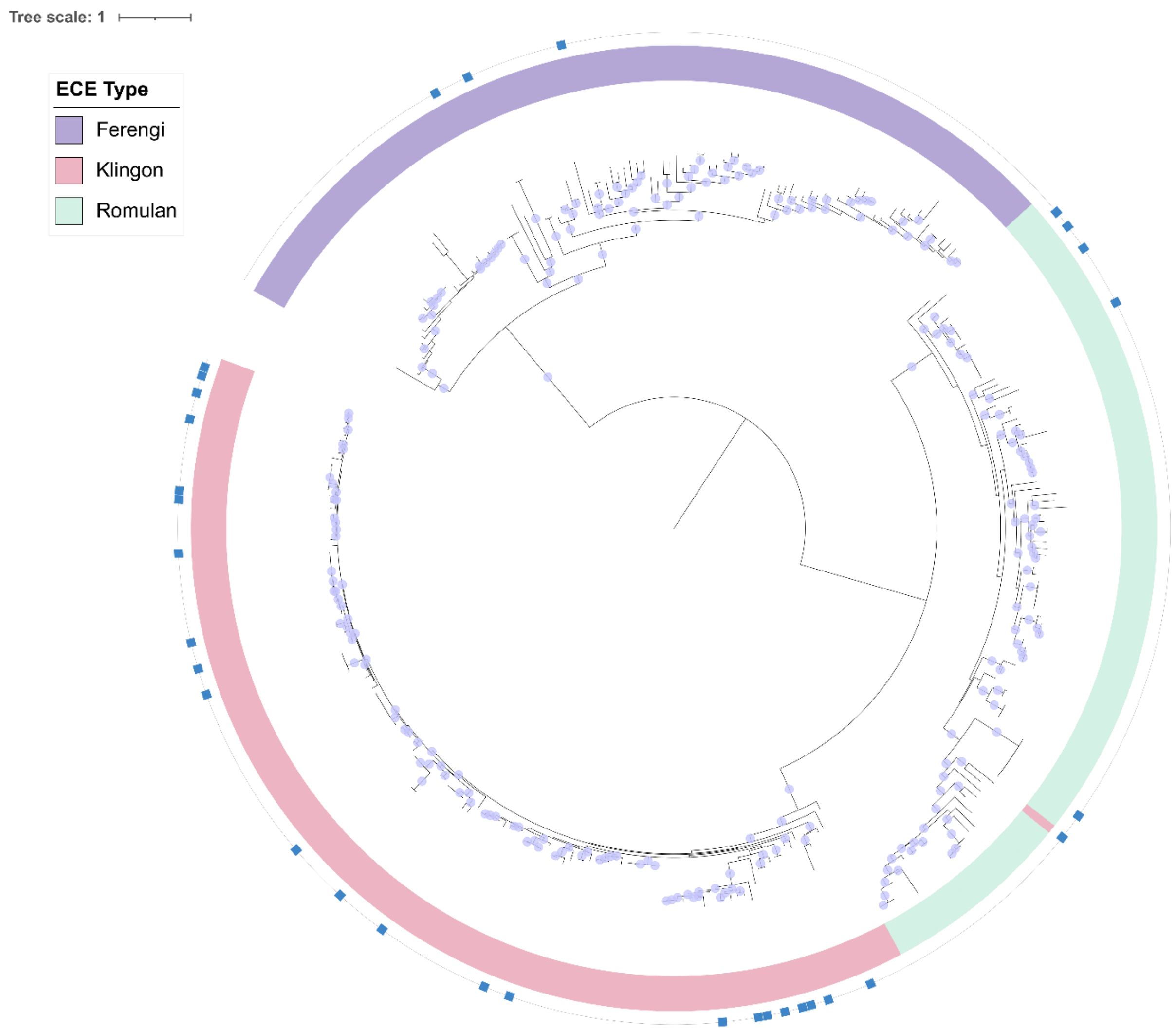


**Supplementary Figure 1: Phylogeny of all curated and partial Klingon, Romulan and Ferengi scaffolds**. Not every FKR genome is included based on presence of marker gene in genome. Bootstrap values (BVs) > 0.70 are marked with lavender dots. Curated genomes are marked on the outer ring with blue squares. The LG+I+R5 substitution model was used for phylogenetic inference. Phylogeny was visualised with iTOL.

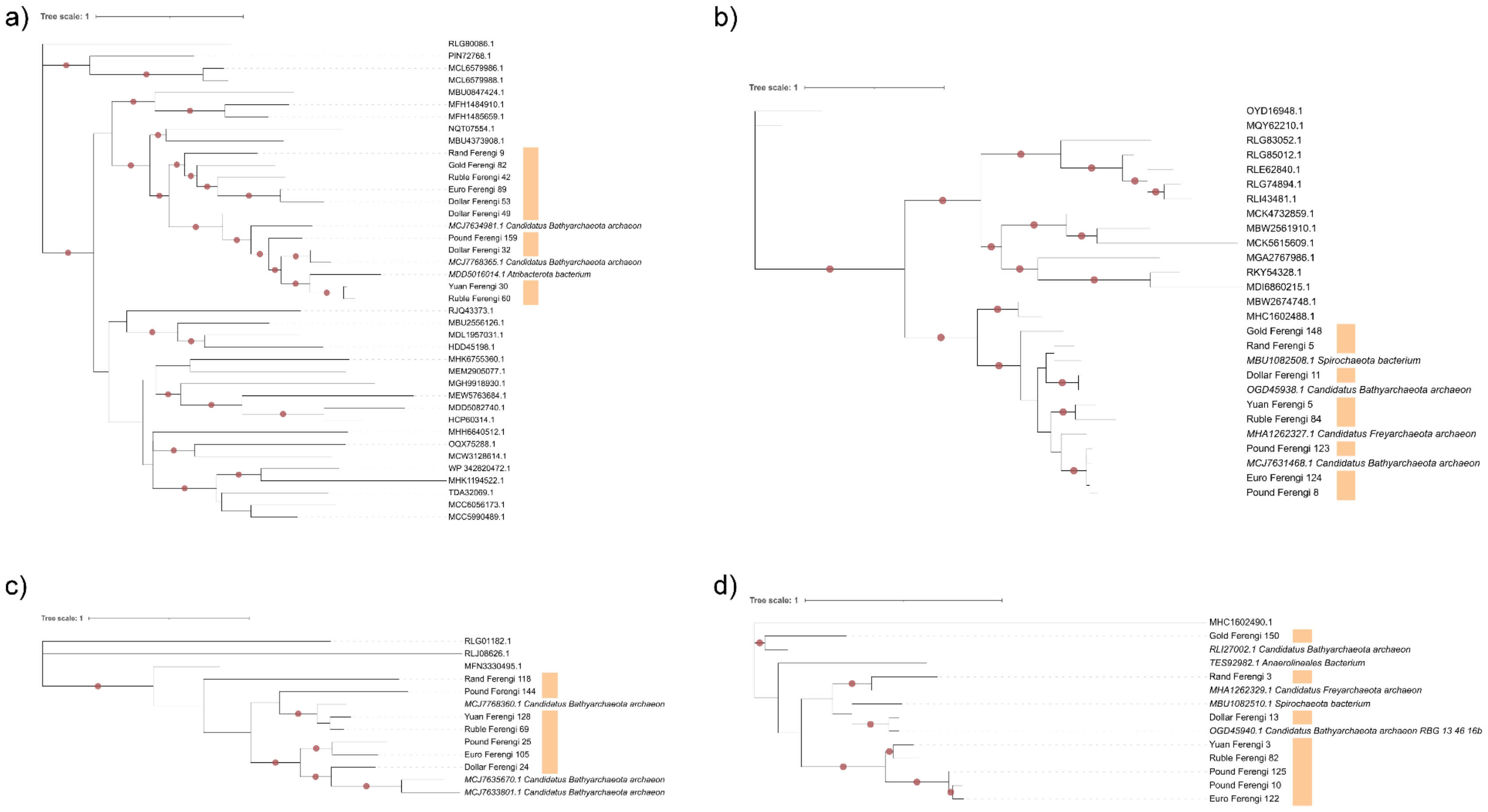


**Supplementary Figure 2: Protein trees Klingon, Romulan and Ferengi proteins with BLASTP matches for host inference.** For all cases, IQTREE was used for phylogenetic inference, and BVs were derived from 1000 ultrafast bootstrap iterations. Ferengi genomes are marked in orange, and BVs > 0.70 are marked in red. Terminal branches within the Ferengi monophyletic lineage were labelled with species name. Subfamilies and substitution models for each tree are as follows, (a) Subfam0542, VT+R3, (b) Subfam 2638, WAG+G4, (c) Subfam3079, WAG+G4, (d) Subfam4970, Q.YEAST+G4. Phylogenies were visualised with iTOL.

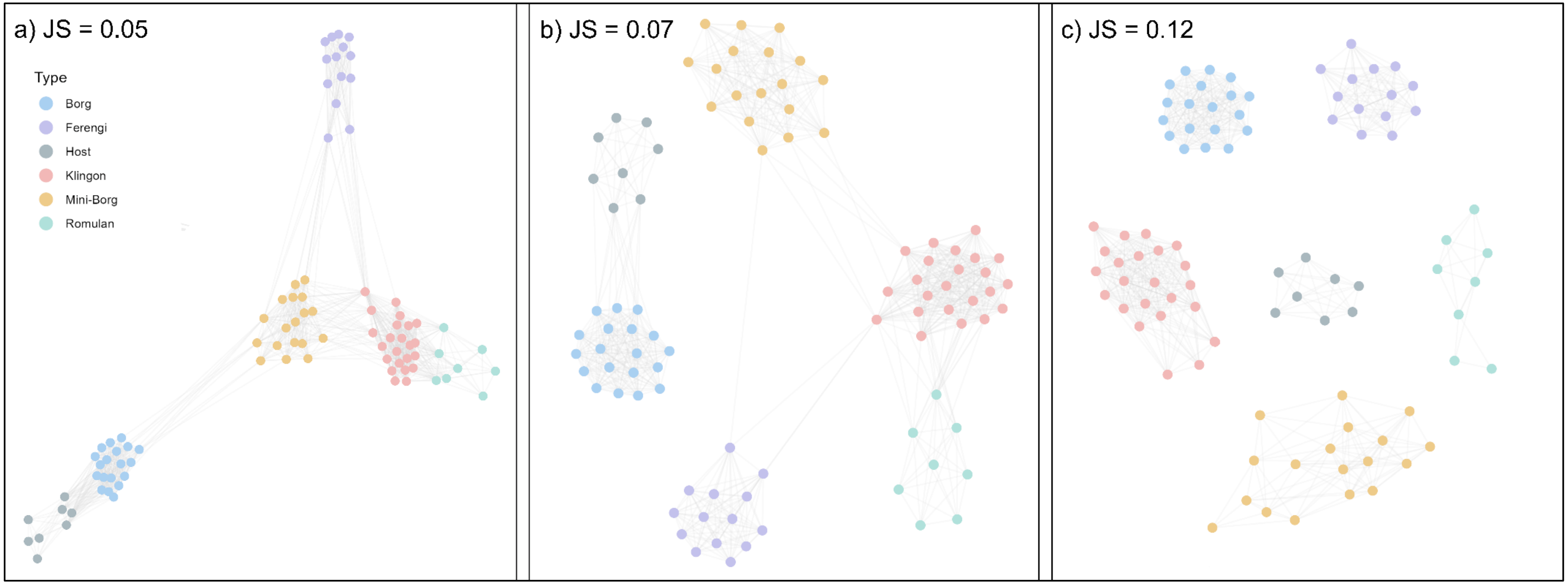


**Supplementary Figure 3: Protein sharing networks of ECE families and potential hosts.** Jaccard similarity (JS) weighting was applied to account for different genome sizes of ECEs and hosts. (a) JS weighting = 0.05, (b) JS weighting = 0.07, (c) = JS weighting = 0.15. R packages “ggplot2”, “ggraph” and “igraph” were used for visualisation.

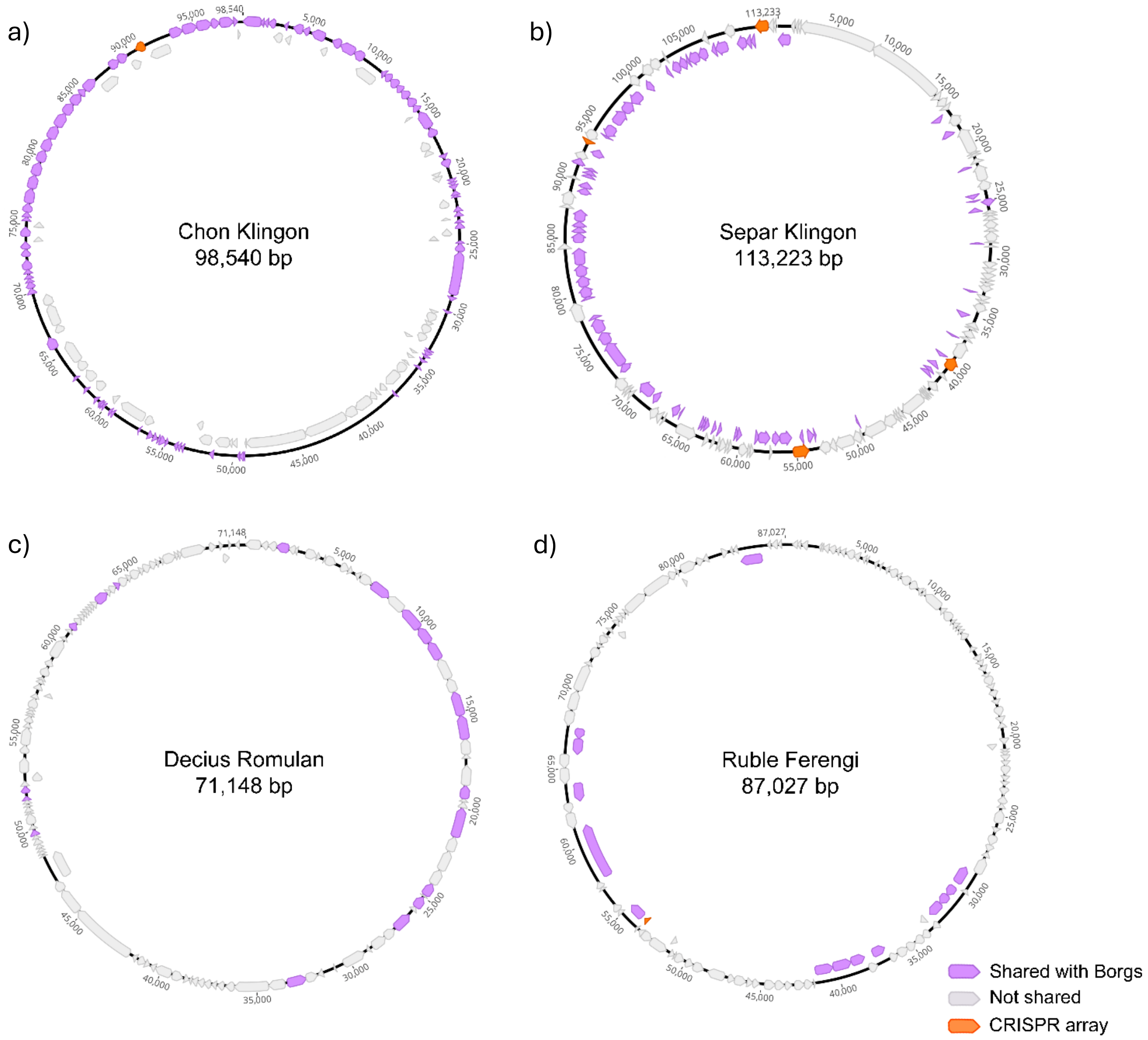


**Supplementary Figure 4: Borg-ECE shared genes have a positional bias in Klingons, Romulans and Ferengi.** Purple annotations mark a gene shared between the ECE family and Borgs in (a) Chon Klingon, (b) Separ Klingon, (c) Decius Romulan and (d) Ruble Ferengi. Shared genes were derived from subfamilies with at least three Borg representatives and three representatives from the respective ECE family. CRISPR arrays are marked with orange annotations. Geneious v2025.2 was used for annotation and visualisation.

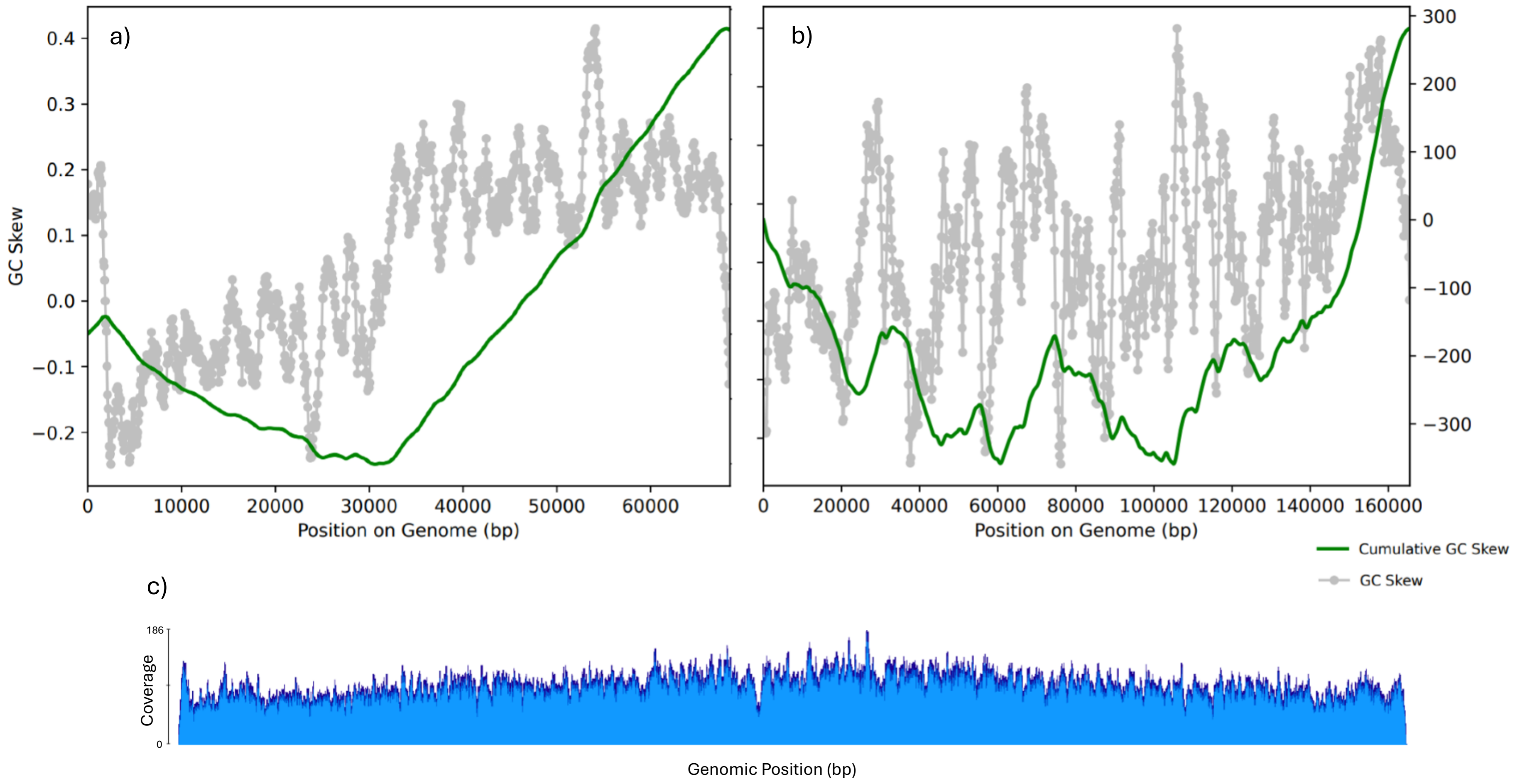


**Supplementary Figure 5: Mapping of FKR genomes according to GC Skew and Coverage.** In plots (a) and (b), the grey line displays the GC skew, and the green line shows cumulative GC skew throughout the genome. (a) GC Skew plot of Dividices Romulan, in which origin and terminus can be determined from GC Skew alone. (b) GC Skew plot of Bas Klingon, in which origin and terminus cannot be confidently determined from GC Skew (c) 105,195 DNA reads mapped with 1% SNP allowance with Geneious Mapper. Origin was adjusted according to coverage peak. Visualisation of the read mapped file was performed with Geneious 2025.2.

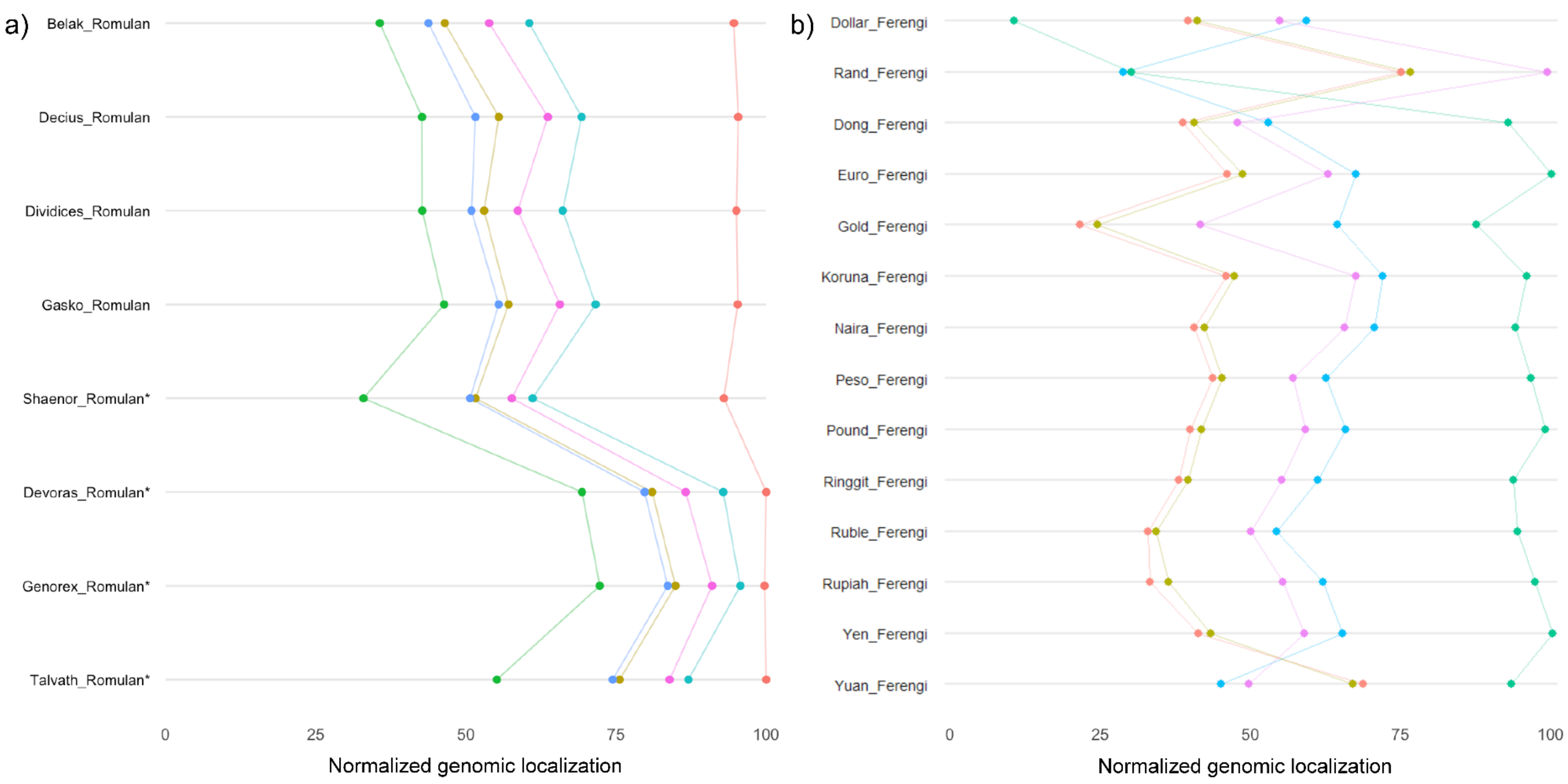


**Supplementary Figure 6: Single copy genes have a mostly conserved order in (a) Romulans and (b) Ferengi.** Romulan-like elements in a) are marked with an asterisk. ECEs' start position has been re-assigned to ensure linearisation of the genomes did not distort the pattern of conserved order.

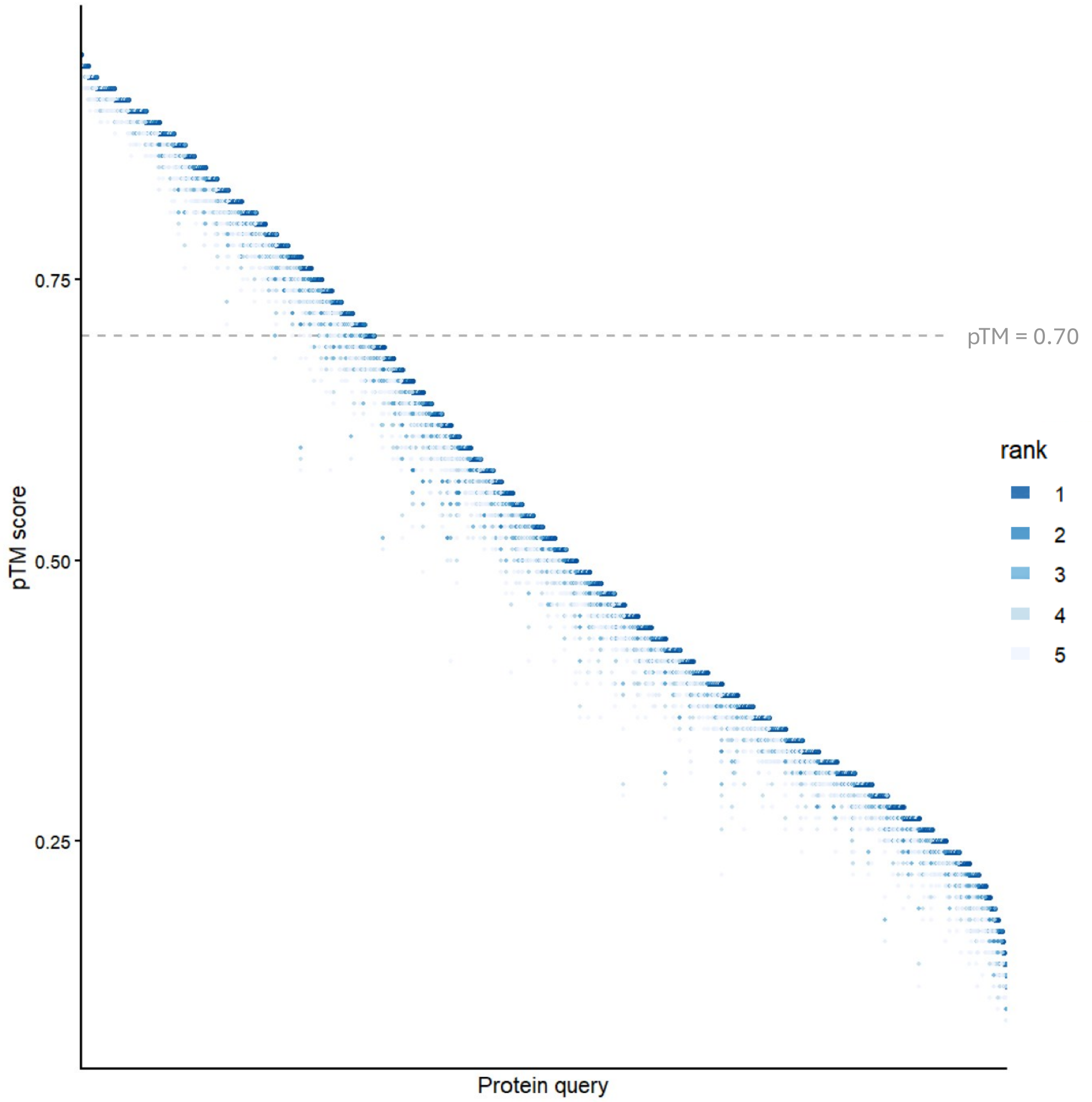


**Supplementary Figure 7: Rainfall plot of pTM values of models 1 - 5 for each ColabFold protein model from the Klingon, Romulan and Ferengi proteins.** The dotted line represents the pTM > 0.70 threshold applied for inclusion in the matrix analysis.

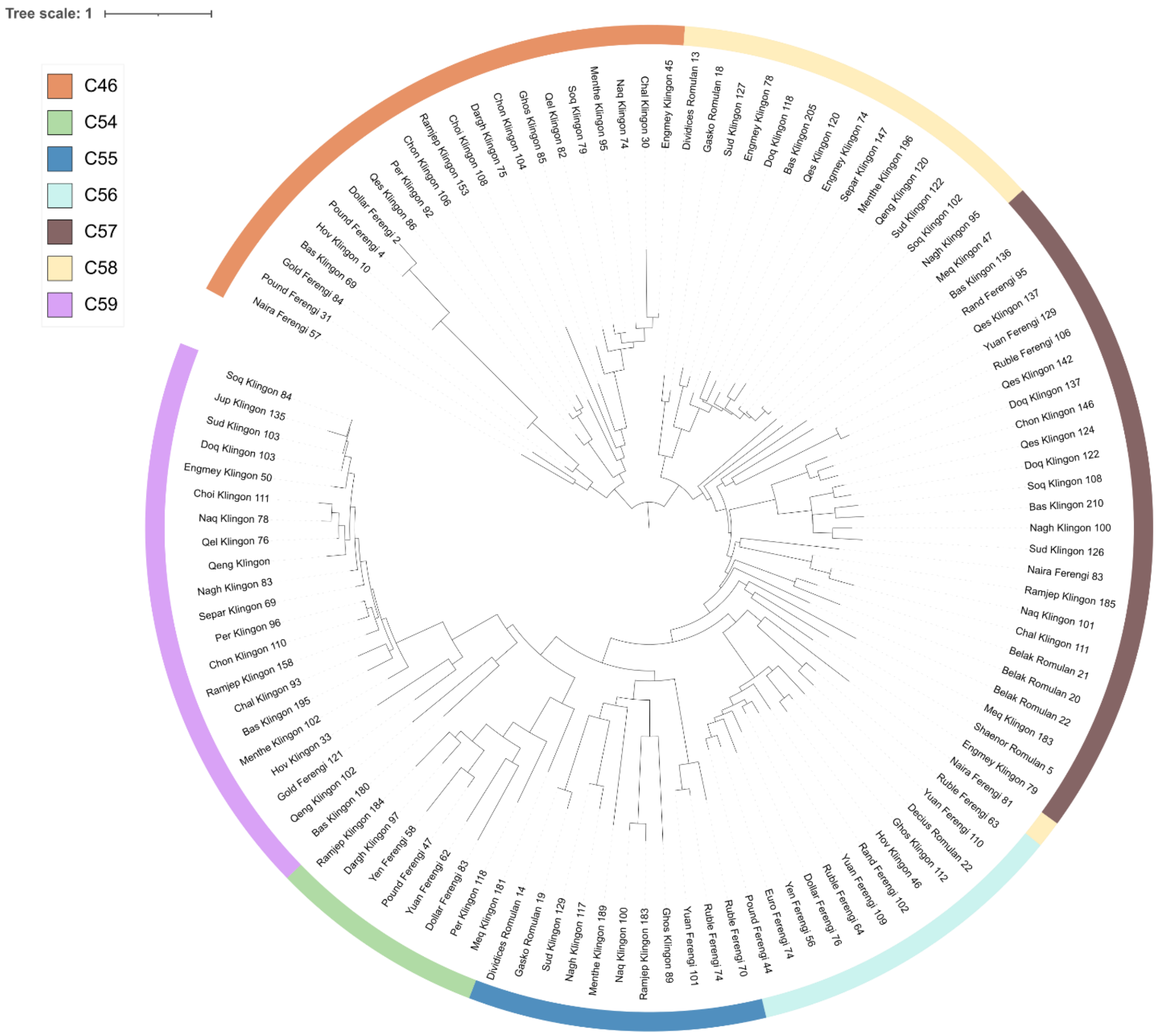


**Supplementary Figure 8: Structure-based phylogeny of FKR-encoded methyltransferase proteins from C46 and C56 - C59.** Cluster (C) assignments are from structural clustering matrix of FKR proteins (Figure 5). Phylogeny was rooted at the base of the C46 lineage based on average similarity to other clusters. Phylogeny was inferred via FoldTree and visualised using iTOL.

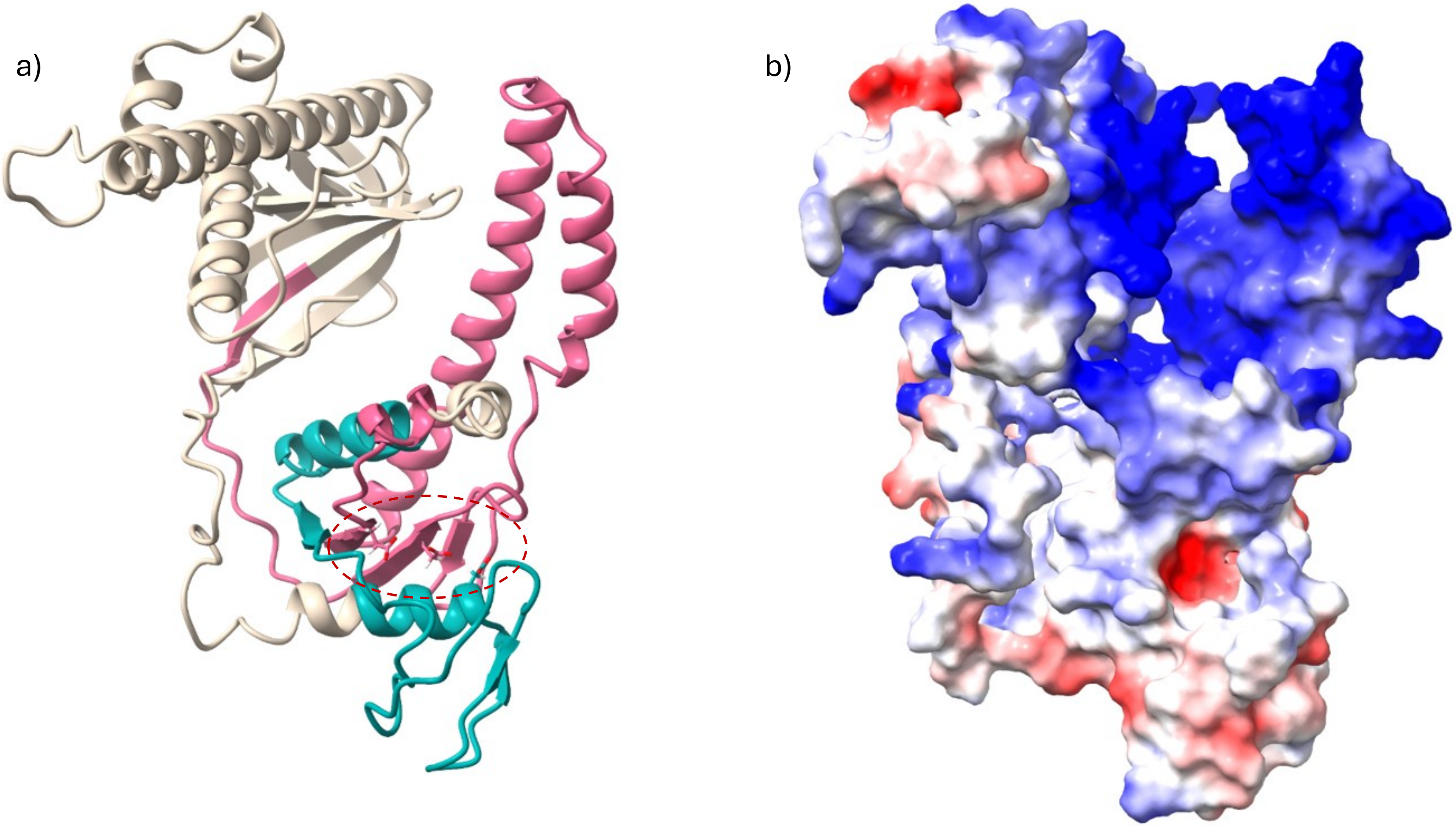


**Supplementary Figure 9: Klingon-encoded nuclease proteins are structurally similar to known TnpB-like nucleases.** (a) In silico protein model of the Ramjep Klingon-encoded RNA-guided endonuclease (Ramjep 88), predicted by Colabfold. DNA binding domain (PF07282) is annotated in teal, transposase domain (PF01385) is annotated in pink. Chains of DED box residues of the RuvC domain are contained within dotted circle. (b) Electrostatic map of Ramjep 88. Proteins visualised with ChimeraX

### List of Supplementary Tables

**Supplementary Table 1:** Sequence-based protein clustering of all FKR, Borg, Mini-Borg and predicted host proteins

**Supplementary Table 2:** Non-FKR genomes in study

**Supplementary Table 3:** Similarity of FKR proteins to other ECEs and predicted hosts

**Supplementary Table 4:** Spacer Matches to *Methanoperedens* in Klingon-encoded CRISPR-like arrays

**Supplementary Table 5:** Relative synonymous codon usage in FKR genomes

**Supplementary Table 6:** Mann-Whitney U test results for codon influence on clustering

**Supplementary Table 7:** GeNomad Statistics

**Supplementary Table 8:** All-against-all comparison of structural similarity of protein models as measured by TM-Score

**Supplementary Table 9:** Methylation Host-ECE linkage analysis

**Supplementary Table 10:** Capsid-like protein multimer analysis

### List of Supplementary Data

Supplementary available via https://doi.org/10.5281/zenodo.23050150

**Supplementary Data 1:** Protein trees used as input for coalescence-derived phylogeny in Figure 2a

**Supplementary Data 2:** ColabFold generated PDB files from structural alignment matrix (Figure 5) large clusters

**Supplementary Data 3:** Amino acid alignment of FKR-encoded NAD-dependent epimerase/dehydratases from cluster 7.

**Supplementary Data 4:** Amino acid alignment of FKR-encoded TnpB proteins and select references from Chen et al (2023) and Xiang et al (2023).
